# ILC1-derived TGFβ1 drives intestinal remodelling

**DOI:** 10.1101/2020.04.20.051805

**Authors:** Geraldine M. Jowett, Michael D. A. Norman, Tracy T. L. Yu, Patricia Rosell Arévalo, Dominique Hoogland, Suzette Lust, Emily Read, Eva Hamrud, Nick J. Walters, Umar Niazi, Matthew Wai Heng Chung, Daniele Marciano, Omer Serhan Omer, Tomasz Zabinski, Davide Danovi, Graham M. Lord, Jöns Hilborn, Nicholas D. Evans, Cécile A. Dreiss, Laurent Bozec, Oommen P. Oommen, Christian D. Lorenz, Ricardo M.P. da Silva, Joana F. Neves, Eileen Gentleman

**Author notes:** These two authors contributed equally; listed in alphabetical order (Michael D. A. Norman, Tracy T. L. Yu). These two authors contributed equally (Joana F. Neves, Eileen Gentleman).

## Abstract

Organoids can shed light on the dynamic interplay between complex tissues and rare cell types within a controlled microenvironment. Here, we developed gut organoid co-cultures with type-1 innate lymphoid cells (ILC1) to dissect the impact of their accumulation in inflamed intestines. We demonstrate for the first time that murine and human ILC1 secrete TGFβ1, driving expansion of CD44v6^+^ epithelial crypts. ILC1 additionally express MMP9 and drive gene signatures indicative of extracellular matrix remodelling. We therefore encapsulated human epithelial-mesenchymal intestinal organoids in MMP-sensitive, synthetic hydrogels designed to form efficient networks at low polymer concentrations. Harnessing this defined system, we demonstrate that ILC1 drive matrix softening and stiffening, which we suggest occurs through balanced matrix degradation and deposition. Our platform enabled us to elucidate previously undescribed interactions between ILC1 and their microenvironment, which suggest that they may exacerbate fibrosis and tumour growth when enriched in inflamed patient tissues.

## Main

Intestinal epithelial cells (IEC)^1^ interact with innate lymphoid cells (ILC)^2^ to form a dynamic barrier between organisms and their environment. Together, they are capable of rapidly responding to danger and damage in an antigen non-specific manner. For instance, type-3 ILC3 secrete Interleukin-22 (IL-22, *Il22*) in response to extracellular pathogens, which promotes anti-microbial peptide secretion and proliferation of Lgr5^+^ CD44^+^ intestinal stem cells^3^. Conversely, type-1 ILC express Interferon-gamma (IFNγ, *Ifng*) in response to intracellular pathogens, and are comprised of circulating natural killer (NK) cells and tissue resident helper-like ILC1 (ILC1), which are considered less cytotoxic than their NK-cell counterparts^4^. Notably, ILC1 accumulate in the inflamed intestines of Inflammatory Bowel Disease (IBD) patients^5^, however the nature of their subset-specific interactions with the epithelium has remained elusive. Understanding the impact of ILC1 enrichment could offer novel avenues for treating this complex disease, which is a pressing issue as only a third of patients respond to gold standard TNFα-blocking biologics^6^.

Teasing apart the role of rare cell populations in multifactorial diseases is challenging, and redundant cytokine signalling pathways *in vivo* can obscure ILC-specific phenotypes. Thus, to explore the impact of ILC1 on IEC we developed a reductionist co-culture system with murine small intestine organoids (SIO)^7^. We unexpectedly found that ILC1-derived TGFβ1 induces p38γ activity to drive epithelial *Cd44v6* expression and SIO proliferation. Pathway analysis of co-culture transcriptomes also predicted ILC1-driven matrisome remodelling, so we developed highly defined PEG-based hydrogels to quantitatively characterize the impact of ILC1 on matrix remodelling in a human iPSC-derived organoid model (HIO)^8^. We not only confirmed that IBD patient-derived ILC1 express *TGFB1* and upregulate CD44v6, but also that they prompt physical changes in the hydrogel via both degradation and production of peri-organoid matrix. Our findings suggest that ILC1 play a role in intestinal remodelling, which could exacerbate IBD-associated comorbidities when enriched in inflamed intestines.

## Results

### ILC1 drive CD44^+^ crypt expansion

To study the impact of ILC1 accumulation on IEC, we established co-cultures of murine SIO and small intestinal lamina propria-derived ILC1 (Fig. 1a-c and Supplementary Fig. 1). ILC1 maintained characteristic KLRG1^-^, RORγt^-^, NK1.1^+^ expression after co-culture (Supplementary Fig. 2, 3a), and expressed *Ifng*, but not *Il22*, matching freshly isolated ILC1 (Fig. 1d). We tuned this system to contain low-levels of IFNγ without driving excess SIO cytotoxicity (Supplementary Fig. 3b,c), and cultured SIO either alone or with ILC1 for 4 days. We then FACS-purified IEC for bulk Smart-seq2 pico-RNAsequencing. ILC1 co-culture significantly increased expression of *Cd44*, a common crypt stem cell marker that can act as a growth factor co-receptor, a transcription factor, or mediate cell surface adhesion^9^ (Fig. 1e, Supplementary Fig. 2, and Supplementary Data Set 1). ILC1 also increased the number of enlarged CD44^+^ crypt buds per organoid (Fig. 1f-h). To explore whether IFNγ drove this effect, we supplemented SIO-only cultures with recombinant IFNγ, but found that it could not account for the increase in epithelial cell numbers and *Cd44* expression (Fig. 1i, j). Moreover, Ingenuity Pathway Analysis (IPA) of the SmartSeq2 dataset did not predict *IFNG* as a dominant signature of ILC1 co-culture (Fig. 1k), suggesting that ILC1 upregulate CD44 through an alternate mechanism.

**Figure 1:**
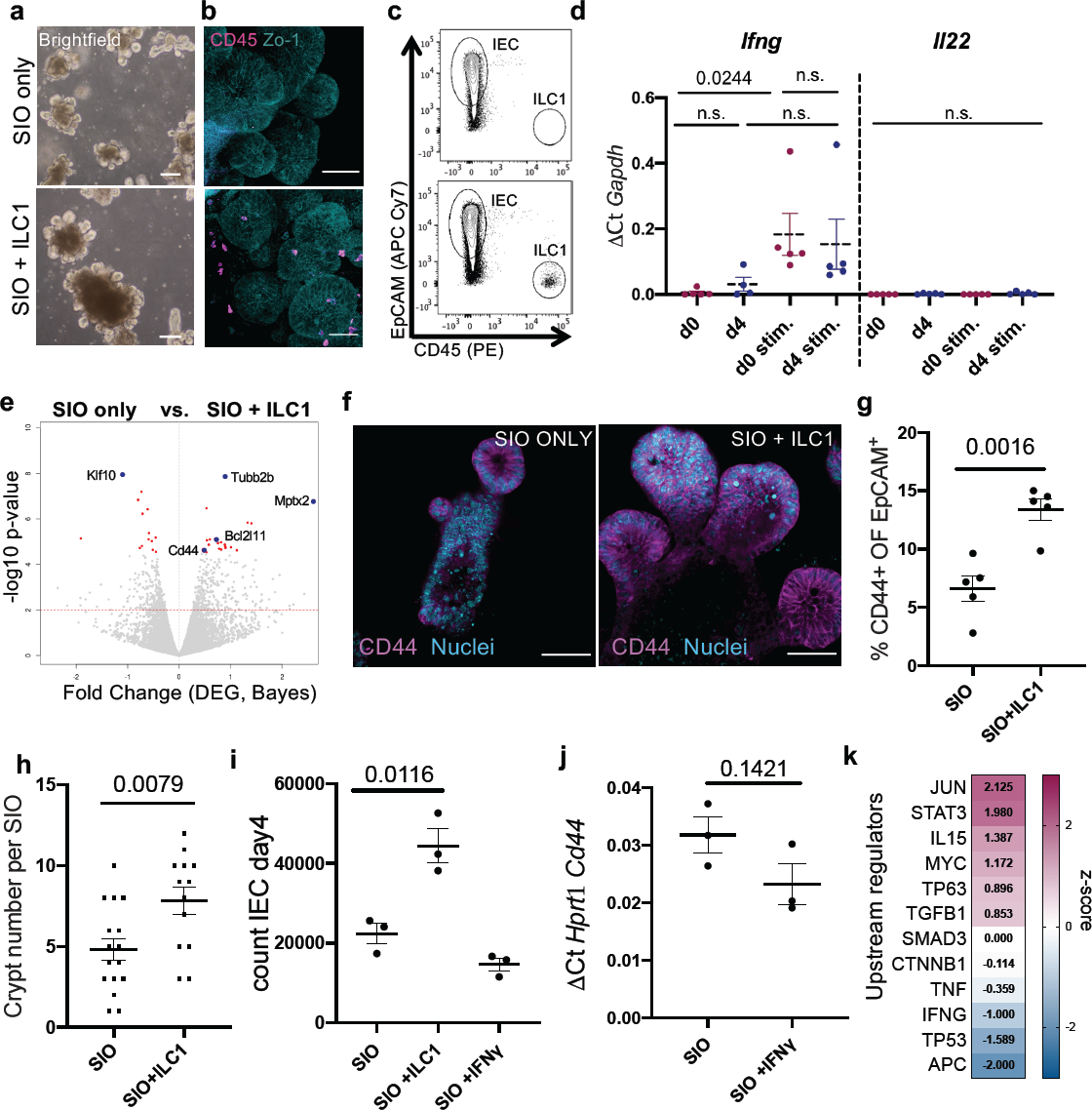
ILC1 impact intestinal organoid gene expression. Representative a. brightfield images, b. confocal images, and c. FACS plots of SIO cultured alone (top) or with ILC1 (bottom) (ILC1 from N=3 mice) d. Expression of *Ifng* and *Il22* in ILC1 pre and post SIO co-culture, with or without 2h stimulation with 10ng/ml PMA & 1*µ*M Ionomycin (stim.). e. Volcano plot (log_10_ p_adj_-value vs. fold change) of differentially expressed genes in pico-RNAsequencing dataset, with significantly upregulated genes of interest highlighted with a blue dot. f. Confocal images of SIO showing CD44^+^ crypts with Lyzozyme1^+^ (Lyz1) Paneth cells (Rep. of N=3). g. Flow quantification of CD44^+^ IEC (%) (N=5). h. quantification of crypt bud number per SIO (N=3, 3-6 SIO per image). i. Number of EpCAM^+^CD45^-^ IEC in SIO after 4 day co-culture alone, with ILC1, or with 0.01ng/ml IFNγ (N=3). j. RTqPCR of IEC *Cd44* expression with or without 0.01ng/ml IFNγ (N=3). k. Ingenuity pathway analysis of selected upstream regulators predicted to be driving expression signatures in ILC1 co-cultures, z-score magenta predicts high activity, blue predicts low activity. d,g,h,i,j show unpaired two-tailed t-test p-values between conditions; error bars S.E.M.; scale bars 50*µ*m.

### ILC1 secrete TGFβ1

As predicted by the IPA upstream regulators (Fig. 1k), we identified increased levels of TGFβ1 in the ILC1 co-culture supernatants (Fig. 2a). Stimulated ILC1 expressed *Tgfb1* before and after co-culture (Fig. 2b), mimicking expression patterns of *Ifng* (Fig. 1d). Conversely, although IECs can upregulate *Tgfb1* in response to microbiome metabolites^10^, IEC expression of *Tgfb1* was negligible both with and without ILC1 co-culture (Fig. 2c). However, SIO in our system did still express TGFβR1 (Fig 2d), and upregulated this receptor in the ILC1 co-cultures (Supplementary Fig. 4), indicating that epithelial cells retained the capacity to respond to exogenous TGFβ1.

**Figure 2:**
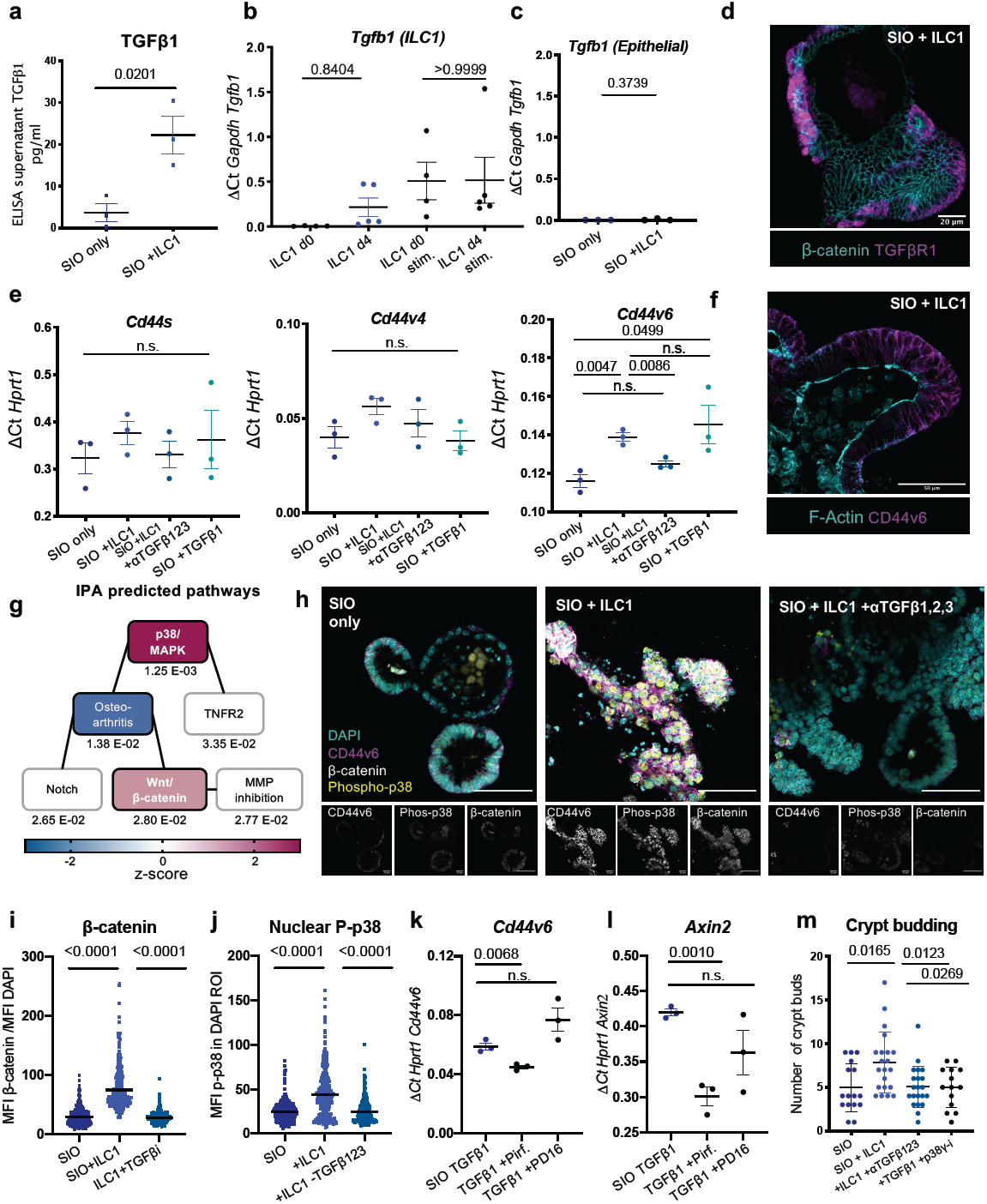
ILC1 impact epithelial crypt gene expression through TGFβ1 secretion. a. ELISA for latent and active TGFβ1 in culture supernatants on day 4 (N=3). b. Expression of *Tgfb1* in primary murine ILC1 before (d0, N=4) or after co-culture (d4, N=5), with or without 2h PMA/Ionomycin activation (stim.). c. Expression of *Tgfb1* in IEC from SIO only or SIO+ILC1 on d4, (N=3, x-axis scaled to 2b). d. Localization of TGFβR1 staining in SIO co-cultured with ILC1 (d4) counterstained for β-catenin (Rep. of N=3). e. RT-qPCR with exon-specific primers for CD44 splice variants s, v4, and v6 (N=3). f. Representative confocal image of CD44v6 localization in d4 SIO+ILC1, counterstained with F-Actin (Rep of N=3 mice). g. IPA of activated canonical pathways. Linked boxes contain overlapping differentially expressed genes (p_adj_ beneath boxes, black outlined boxes p<0.05; magenta-midnight coloring represents z-score). h. Expression patterns of CD44v6, phosphorylated p38, β-catenin and DAPI nuclei in SIO alone, with ILC1, or with ILC1 and 1ng/ml TGFβ1,2,3-neutralization (Rep of N=3 mice). i. Quantification of phosphorylated p38 in the DAPI^+^ region of the experiment in (h) (N=3, each dot represents one nucleus) j. Quantification of β-catenin accumulation in IEC of experiment in (h), normalized to DAPI intensity (N=3, each dot represents one cell). k. *Cd44v6* expression and l. *Axin2 expression* after 2day culture of SIO with TGFβ1, with TGFβ1 and Pirfenidone (5*µ*m) or TGFβ1 and PD16 (3*µ*m). m. Quantification of crypt budding after 4 day co-culture (OneWay ANOVA and Tukey test N=3, 5-7 SIO per condition). a,b,c,e,k,l Two-tailed unpaired t-test, error bars show S.E.M.; i,j,m, OneWay ANOVA and Tukey test, error bars S.E.M. Scale bars indicated in overlays.

We next investigated whether TGFβ1 accounted for CD44 upregulation. First, we established that the phenotype was not contact dependent (Supplementary Fig. 5). We then distinguished between common splice isoforms of CD44 using intron-specific primers^11^ and found that ILC1 co-culture upregulated CD44 variant 6 (*Cd44v6*) specifically, and could be inhibited by TGFβ1,2,3 neutralizing antibody and upregulated by adding recombinant TGFβ1 to SIO-only cultures (Fig. 2e). Importantly, TGFβ1,2,3 inhibition did not adversely impact ILC1 numbers (Supplementary Fig. 6). CD44v6 protein was ubiquitously distributed across the basolateral membrane of the SIO crypt in co-cultures (Fig. 2f), and did not appear to concentrate in specific IEC subsets. TGFβ1-induced expression of CD44v6 has been described in fibrotic lung fibroblasts^12^, however this is to our knowledge the first description of such a connection in the intestinal epithelium.

CD44 engages in a positive feedback loop with Wnt/β-catenin. Indeed, it is a downstream target of β-catenin, and clusters with Lrp6 to potentiate Wnt signalling^13^. Moverover, IPA predicted significant increases in both p38/MAPK and Wnt/β-catenin signalling in SIO co-cultured with ILC1 (Fig. 2g). We observed accumulation of β-catenin in ILC1 co-cultures (Fig 2h), and increased expression of β-catenin-targets *Ascl2* and *Axin2* (Supplementary Fig. 7a). β-catenin accumulation co-localized with CD44v6^+^ cells (Supplementary Fig. 7b), and was reversible by TGFβ1,2,3 neutralization (Fig. 2i), hinting at a potential mechanistic link. We first hypothesized that increased crypt size in ILC1 co-cultures could be driven by CD44-mediated modulation of Wnt-signalling, however despite a trending bias toward increased expression of stem cell crypt over mature enterocyte markers, differences in subset-specific genes were not statistically significant (Supplementary Fig. 8). Instead, IEC that upregulated CD44v6 and β-catenin also showed a dramatic increase in phosphorylated p38 signal (Fig. 2h), which was equally upregulated by ILC1 co-culture and downregulated through TGFβ1,2,3 neutralization (Fig. 2j). This kinase exists in multiple isoforms, and while p38α/β regulates apoptosis, p38γ promotes proliferation. To investigate which isoform was active in our co-cultures, we used p38α/β-inhibitor PD169316 and p38γ-inhibitor (PD16) Pirfenidone, a drug approved for the treatment of pulmonary fibrosis^14^. These soluble inhibitors impacted ILC1 phenotypes (Supplementary Fig. 9a), so we mimicked ILC1 co-culture through addition of recombinant TGFβ1. SIO cultured with Pirfenidone, but not PD16, significantly and specifically downregulated *Cd44v6* (Fig. 2k, Supplementary Fig. 9b) and *Axin2* (Fig. 2l), and reversed crypt budding similarly to TGFβ1,2,3 neutralization (Fig. 2m). It is reported that p38γ phosphorylates the Ser605 residue of β-catenin, thus stabilizing it and driving inflammation-associated intestinal tumorigenesis^15^. This suggests that p38γ activity likely acts downstream of TGFβ1 and upstream of β-catenin and CD44v6 upregulation, which could promote IEC subtype non-specific proliferation and organoid growth through a positive feedback loop.

### IBD-patient ILC1 upregulate intestinal CD44v6

To add translational value to these data, we isolated human intestinal lamina propria ILC1 (hILC1) from IBD patient biopsies (Supplementary Fig. 10), and established co-cultures with human gut organoids (Supplementary Fig. 11, 12). Epithelial-only biopsy-derived enteroids^16^ closely mimic SIO, but as these maintain epigenetic signatures of their donors^17^, they offer no control over patients’ genetic background or exposure to environmental stressors, diet, or drugs. Conversely, differentiation^18^ and maturation^19^ of human iPSC-derived intestinal organoids (HIO) provides greater control over genetics and environment, which are necessary for modelling a multifactorial disease. Following 7-day co-culture with HIO, hILC1 maintained their phenotypic response to activation, upregulating *IFNG* but not *IL22* (Fig. 3a). In this system, hILC1 co-culture increased basolateral CD44v6 expression in HIO (Fig. 3b), which was recapitulated through addition of recombinant TGFβ1 (Supplementary Fig. 13). Strikingly, the increase in *CD44v6* expression was only statistically significant in cultures with hILC1 derived from sites of active inflammation (Fig. 3c). These biopsies also yielded the reported increased ratio of hILC1 relative to other ILC subtypes^5^ (Fig. 3d, Supplementary Fig. 10). hILC1 expressed *TGFB1* before (Supplementary Fig. 14) and after co-culture with HIO (Fig. 3e), but expression of *TGFB1* did not differ significantly between inflamed and uninflamed samples. However, the number of hILC1 in cultures from inflamed tissues increased significantly in co-culture, while those from uninflamed biopsies did not (Fig. 3f). It is therefore probable that the differential increase in *CD44v6* in inflamed co-cultures results from the increased number of hILC1 in these conditions. These observations confirm that inflamed-biopsy-derived ILC1 are most appropriate for translating our murine data. Moreover, as HIO phenotypes and genotypes were consistent between conditions, they suggest that the inflamed IBD environment leaves a lasting imprint on hILC1, which is maintained *in vitro* in the absence of other immune cells, genetic abnormalities, environmental damage, or a dysregulated microbiome.

**Figure 3:**
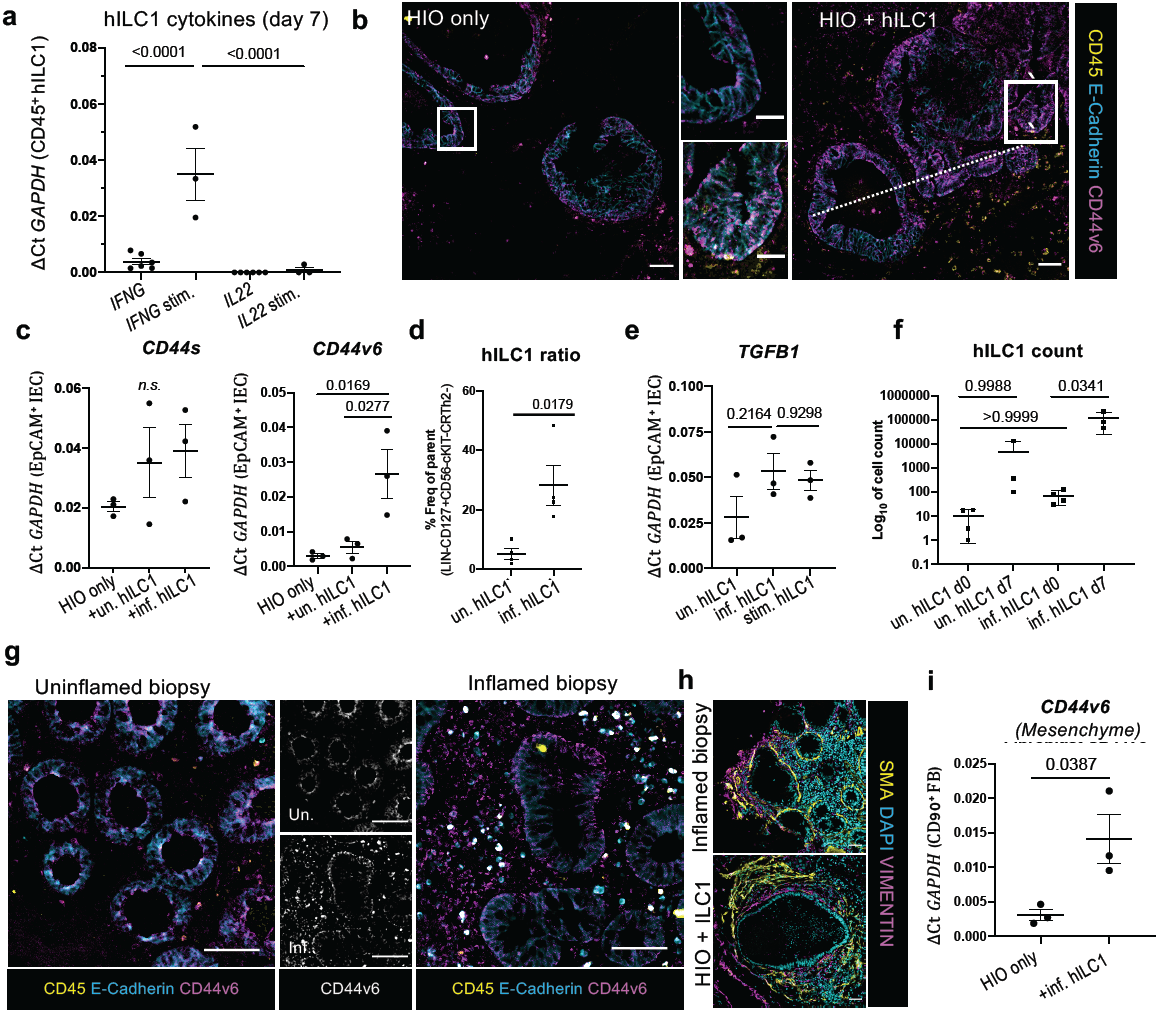
Human ILC1 drive CD44v6 expression in HIO. a. Expression of *IFNG* and *IL22* in biopsy-derived hILC1 after 7 day co-culture with HIO with (N=3) or without (N=6) 2h PMA/Ionomycin stimulation. b. E-cadherin, CD44v6, and CD45 staining of HIO cultures with or without hILC1 (Scale 50*µ*m), white box indicates magnified crypt (Scale 20*µ*m, Rep. of N=3). c. Expression of *CD44s* and *CD44v6* in FACS-purified IEC from HIO only, hILC1 co-culture, from inflamed (inf.) or uninflamed (un.) samples (N=3 patients per condition). d. Proportion of hILC1 relative to other Lineage^-^CD127^+^ ILC subtypes prior to co-culture (N=4). e. Relative expression of *TGFB1* in hILC1 from inf., un., or stim. samples after 7 day co-culture with HIO. f. Log_10_ cell count of hILC1 before (d0, N=4) and after (d7, N=3) co-culture from un. and inf. biopsies (Rep. of N=3). g. Immunohistochemistry from biopsies from IBD patients with (right) or without (left) active inflammation. CD45 lymphocytes also express E-cadherin and CD44v6. h. Example of comparable SMA+ myofibroblast and VIMENTIN+ fibroblast organization around the epithelium in an inflamed patient biopsy (left) and in HIO+ILC1 co-cultures (right). i. Relative expression of *CD44v6* in EpCAM^-^CD45^-^CD90^+^ fibroblasts (FB) purified from HIO after 7 day culture with or without inflamed hILC1. a,c,e,f OneWay ANOVA with Tukey’s test, error bars S.E.M; d,i, unpaired two-tailed t-test, error bars S.E.M.. Scale bars 50*µ*m.

To assess the clinical relevance of our system, we performed immunohistochemistry on inflamed and uninflamed intestinal biopsies. We observed an increase in epithelial CD44v6 expression along the basolateral junctions of enlarged crypts in inflamed tissues. This underscored that co-cultures of SIO+ILC1 and HIO+hILC1 (from patients with active inflammation) both predicted and recapitulated CD44v6 upregulation in inflamed IBD tissues. However, we also noticed CD44v6 expression beyond the epithelial compartment, in both CD45^+^ lymphocytes and in basal lamina fibroblasts (Fig. 3g). Since inflamed tissues are infiltrated by many different immune cells, we could not determine whether the mesenchymal upregulation was related to hILC1 accumulation, and therefore returned to the HIO model. HIO co-develop with organized layers of mesenchymal fibroblasts, closely mimicking the ECM environment of the native intestine (Fig. 3h). We found that HIO fibroblasts expressed significantly more *CD44v6* after co-culture with hILC1 from inflamed tissues (Fig. 3i), suggesting a causal link between hILC1 and mesenchymal remodelling. Since TGFb1 is a master regulator of fibrosis, and pathological matrix remodelling is a hallmark of IBD^20^, this merited further investigation.

### Modular synthetic hydrogels provide quantitative assessment of matrix remodelling

The responsiveness of fibroblasts to hILC1 piqued our interest, as Gene Set Enrichment Analysis (GSEA) of the murine SIO dataset had revealed significant enrichment of ECM-remodelling genes in co-culture (Supplementary Fig. 15). We also frequently observed degradation of Matrigel in ILC1 co-cultures, which was reversible through MMP-inhibition (Supplementary Fig. 16a-d). Moreover, we found that ILC1 express gelatinase MMP9, a biomarker for IBD^21^ (Supplementary Fig. 16e,f). Until this point, experiments were conducted by resuspending cultures in 3D mouse sarcoma-derived Matrigel. This laminin-rich gel could mask matrix deposition by fibroblasts, and while it is degradable by native enzymes, the manufacturer adds proprietary concentrations of undefined MMP inhibitors^22^, precluding experiments that require precise control over and quantification of matrix remodelling.

To appropriately address this question, we required a highly defined 3D system with physical properties akin to the native intestine, but whose degradability could be independently modulated. PEG-based hydrogels with suitable stiffness have been reported, but require cross-linking by transglutaminase Factor XIIIa^23^ which is known to crosslink ECM components like fibronectin. Fully synthetic hydrogels in which homo-bifunctional peptides (A_2_) act as crosslinkers of 4- or 8-arm PEGs (B_4_/B_8_) have also been described; however, when crosslinkers bear two identical functional groups that react indiscriminately towards the chain-end of any PEG arm, primary (1°) loop formation^24^ can impact network connectivity. This is critical when forming soft, tissue-like hydrogels which require low polymer concentrations, resulting in slow and inefficient network formation in which organoids reach the tissue culture plastic beneath the hydrogel prior to 3D gelation^25^.

A_4_+B_4_ hydrogel designs that avoid 1° looping could yield more effectively cross-linked networks than A2+B4 systems^26^ (Fig. 4a). To explore if this held true at low polymer concentrations, we carried out molecular dynamics (MD) simulations, using a coarse grain approach. MD uses classical laws of mechanics to provide insight into probable molecular arrangements within a material. Simulations showed that A_4_+B_4_ designs facilitated the formation of more network-forming cross-links than A_2_+B_4_ designs, in which ∼25% of cross-links were 1° loops (Fig. 4b and Supplementary Figs. 17 and 18).

To create an A_4_+B_4_ design, we formed hydrogels using two sequential click reactions in which all peptides acted as cross-linkers (Fig. 4c and Supplementary Fig. 19). First, an amine at peptides’ N-terminal was reacted with PEG-4NPC (A_4_), yielding PEG-peptide conjugates (conjugation efficiency 80-100%). Hydrogels were then formed through a Michael addition between a C-terminal free thiol on the unconjugated peptide arm with the end-terminus of PEG-4VS (B_4_) (>90% efficiency) (Supplementary Fig. 20). A_4_+B_4_ hydrogels had more effectively cross-linked networks with lower swelling ratios (Fig. 4d) that were both stiffer and behaved more elastically (Fig. 4e and Supplementary Fig. 21) than A_2_+B_4_ designs formed using homo-bifunctional peptides. Moreover, although A_4_+B_4_ hydrogels abandoned standard pendant presentations of adhesive ligands, human mesenchymal stromal cells could still adhere to their surfaces (Supplementary Fig. 22). A_4_+B_4_ hydrogels’ Young’s modulus (*E*) could be varied by modulating polymer concentration (Fig. 4f) to achieve values for *E* similar to that of normal human intestinal tissue (750-1250Pa)^27^, and were susceptible to degradation by MMP9 (Fig. 4g). Taken together, this suggests suitable properties to explore HIO matrix remodelling.

### Human ILC1 drive matrix remodelling

Equipped with an appropriate culture system, we found that HIO encapsulated in degradable, non-degradable, and intermediately degradable (IM-DEG, 45% MMP-cleavable peptides) hydrogels were viable (Fig. 4h, Supplementary Fig. 23), and maintained their characteristic phenotype, similarly to Matrigel (Fig. 4 i,j; Supplementary Fig. 24a,b). Moreover, HIO were capable of depositing native extracellular matrix in this system, especially in more degradable gels (Supplementary Fig. 24 c-e). Next, we harnessed this system to quantitatively dissect the impact of hILC1 on the physical properties of HIO-hydrogel cultures. First, we used atomic force microscopy (AFM)-based indentation to map cell-mediated changes in stiffness. Since phenotypically irrelevant differences in mechanical properties between conditions could come from the physical presence of hILC1 within the gel, we opted to surround the HIO-laden hydrogels with ancillary hILC1 from inflamed biopsies (aILC1), keeping the composition of the microenvironment that we mapped constant (Fig. 5a). We collected force-distance measurements using a bead-functionalized cantilever (Supplementary Fig. 25), and observed increased heterogeneity of *E* across maps with aILC1 (Fig. 5b). Indeed, we saw a significant difference in variance of *E* induced by co-culture with aILC1 (F=0.0004; p_K-S_=0.0011)(Fig. 5c). Since aILC1 appeared to induce both stiffening and softening of the matrix, while median *E* remained comparable between samples, this could suggest a balance between cell-mediated matrix production and degradation.

**Figure 5:**
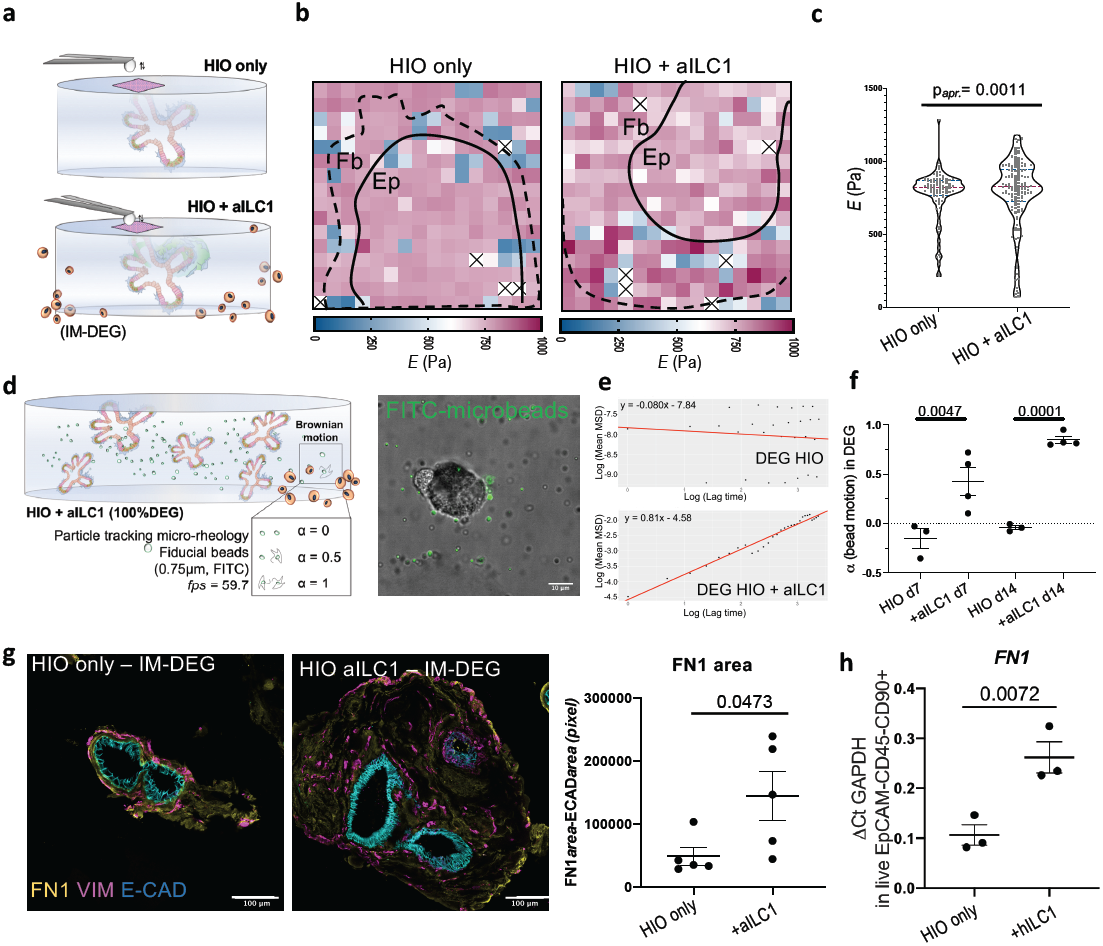
Ancillary ILC1 drive HIO matrix remodelling in synthetic hydrogels. a. Schematic of AFM-based stiffness mapping strategy of HIO in IM-DEG gels, wherein the gel content (HIO) remains constant for measurement, as ancillary hILC1 from inflamed patient tissues surround the gel. b. Representative 150*µ*m x150*µ*m stiffness maps (Pa) of HIO-laden IM-DEG gel without (left) or with (right) aILC1, showing approximate outline of epithelial layer (Ep) and surrounding fibroblast region (Fb) based on brightfield images (Supplementary Fig. 25). White/x squares denote omitted measurements that failed to meet QC standards, Median *E*_HIO_=790.9, Median *E*_HIO+aILC1_=779.9. c. Violin plots summarize measurements of Young’s modulus (*E*, Pa) on HIO-laden IM-DEG gels measured directly above organoids with or without aILC1. F test comparing variances showed a loss of normal distribution in aILC1 culture (p(F)=0.0004), therefore non-parametric Kolmogorov-Smirnov test was used (Approximate p=0.0011, Kolomogorov-Smirnov D=0.2525; 50-70 measurements per force map, N=3). d. Schematic of microrheology strategy (left) and representative confocal image of co-encapsulation (right), wherein fiducial FITC beads are evenly distributed within the HIO-laden fully degradable gel (scale bar 10*µ*m). α is an indicator of bead motion (1=Brownian motion, 0=immobile, α transitions from 0 to 1 as the local hydrogel undergoes a gel-sol transition). e. Representative plots generated in R showing the logarithmic slope of the mean-squared displacement of beads. f. α for HIO encapsulated in 100% DEG gels after 7 and 14 days with or without aILC1 (N=4). g. Representative staining of Fibronectin1 deposition and Vimentin+ fibroblasts in HIO with or without aILC1 (max projection 10 z-stacks) with quantification of FN1+ area normalized to ECAD+ area. h. Expression of *FN1* in EpCAM-CD45-CD90+ FB after 7 day culture with or without inflamed hILC1 (N=3 per condition). f,g, unpaired two-tailed student t-test, h. One-Way ANOVA with Tukey’s test; Error bars S.E.M; scale bars 100*µ*m.

To ensure that aILC1 had the same capacity to degrade HIO-laden hydrogels as in Matrigel, we next performed multiple particle tracking microrheology (Fig. 5d) by monitoring the Brownian motion of fluorescent fiducial beads distributed within the hydrogel. Beads are capable of moving within fully degradable hydrogels when enzyme-mediated degradation causes a sufficient portion of their local environment to undergo a gel-sol transition, prompting the logarithmic slope of a bead’s mean-squared displacement, α, to transition from 0 to 1^28^ (Fig. 5e). After 7 days, aILC1 co-culture resulted in significantly increased α relative to HIO-only controls (Fig. 5f), with near complete degradation after 2 weeks (α=0.847).

As ILC1’s ability to drive degradation of an MMP-sensitive hydrogel could account for the matrix softening recorded by AFM, we next assessed how ILC1 might contribute to the stiffer regions observed in fibroblast-rich, peri-organoid regions. We observed that aILC1 drove a significant increase in the area of peri-organoid FN1 deposition (Fig. 5g, Supplementary Fig. 26a,b). This phenotype was recapitulated in Matrigel, where aILC1 increased expression of *FN1* and *COL1a1* in HIO-fibroblasts (Fig. 5h, Supplementary Fig. 27a), and increased FN1 deposition, similarly to exogenous treatment with TGFβ1(Supplementary Fig. 27b). Specific upregulation of Fibronectin1 is consistent with non-canonical, SMAD4-independent TGFβ1 signalling via Jun/p38, which drives *FN1* expression^29^. We therefore suggest that a balance between ILC1-mediated degradation and ILC1-induced mesenchymal ECM deposition could account for the quantitative difference in coefficient of variance captured in the AFM stiffness maps. In summary, our A_4_+B_4_ defined hydrogel allowed us to conclusively assess that hILC1 drive intestinal matrix remodelling.

## Discussion

Here, we identified ILC1 as an important new source of TGFβ1, which promotes proliferation of CD44v6^+^ epithelial crypts through non-canonical p38γ phosphorylation (Fig. 6a). This is in line with a recently published RNA-sequencing dataset of human ILC1, which showed increased expression of *TGFB1* and *MMP9* in patients with an acute risk of myocardial infarction relative to healthy controls^30^. ILC1 derived from IBD patients with active inflammation also upregulate CD44v6 in both epithelial and mesenchymal cells. To more thoroughly investigate the role of hILC1 in matrix remodelling emerging from our data, we then developed a highly defined synthetic hydrogel system, which allowed us to quantify hILC1-mediated matrix degradation and stiffening.

**Figure 6:**
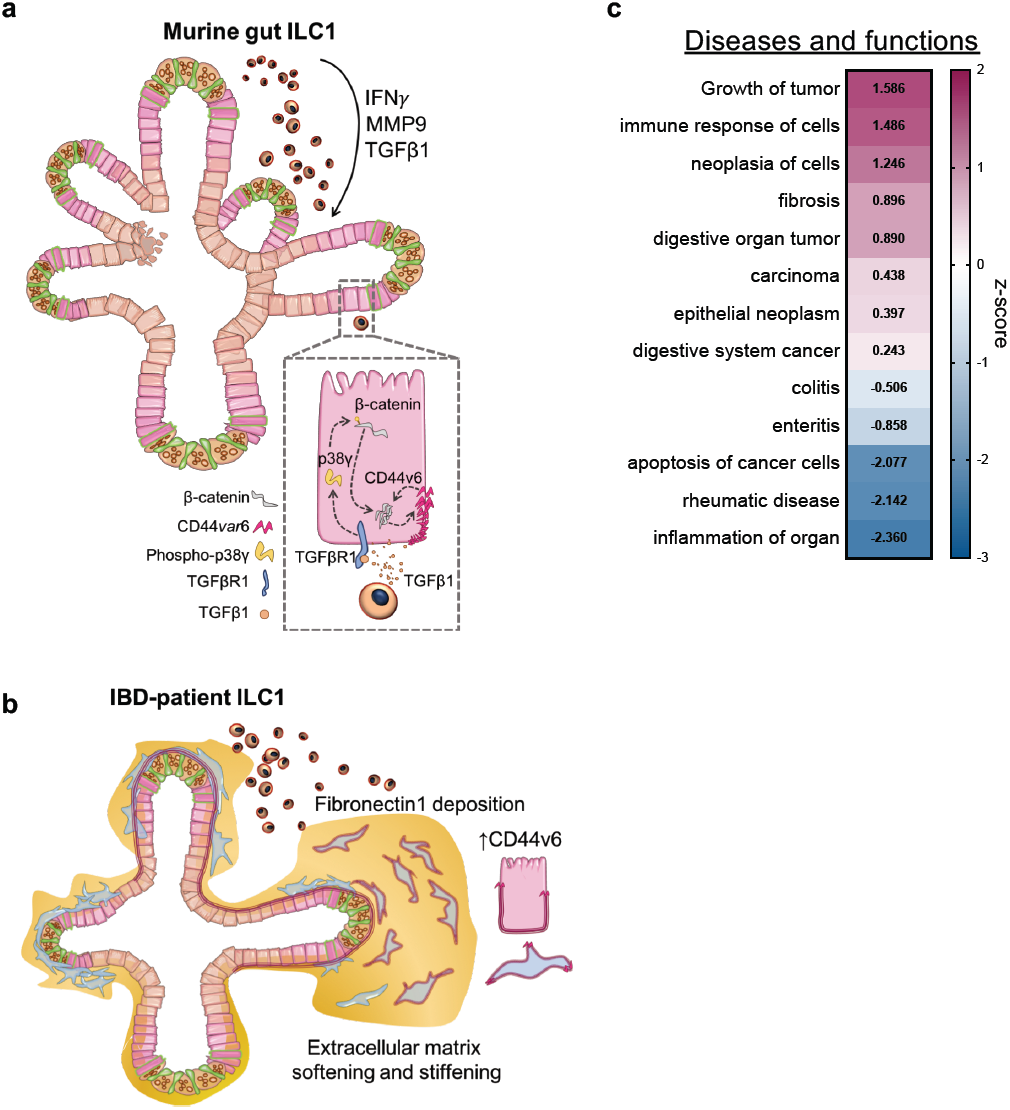
Overview of proposed impact of ILC1 on gut organoids. a. Murine ILC1 drive epithelial crypt budding in small intestine organoids through TGFβ1-induced phosphorylation of p38γ, which drives β-catenin accumulation and expression of *Axin2* and *CD44v6*. We propose that CD44v6 and β-catenin might engage in a positive feedback loop, driving epithelial subtype-non-specific proliferation. b. IBD-patient derived hILC1 express *TGFB1* and *MMP9*. ILC1 isolated from from tissues with active inflammation drive expression of both epithelial and mesenchymal CD44v6 in HIO. Moreover, these patient ILC1 drive increased deposition Fibronectin1 and MMP-mediated matrix degradation, resulting in a balance of matrix softening and stiffening. c. IPA of the murine RNA-sequencing dataset showing cumulative gene enrichment (z-score) in SIO co-cultured with ILC1 indicative of activation (magenta) and inhibition (blue) of selected gastrointestinal and inflammatory diseases and function (*N*=3).

Speculation about the impact of TGFβ1 in the context of the gastro-intestinal immune system is a complex task, as the microbiome, the enteric nervous system, and other immune cells differentially respond to this pleiotropic cytokine. For instance, while TGFβ1 is a master regulator of fibrosis in fibroblasts^31^, it is anti-inflammatory in the adaptive immune system^32^, and can regulate plasticity between the ILC subsets^33^. We observed expression of TGFβ1 in tandem with IFN*γ*, suggesting that these cytokines may act in concert, and highlighting the importance of our dataset being derived from co-cultures with ILC1, not recombinant cytokines.

IPA analysis can potentially address the function of p38γ-induced CD44v6 activity, which suggested a decrease in inflammatory phenotypes and an increase in epithelial gene signatures, consistent with tumour growth and fibrosis (Fig. 6b). This fits with the pathogenic association of this variant, as CD44v6 exacerbates aggressive ovarian cancer by driving β-catenin expression^34^ and drives intestinal cancer initiation^35^, progression^9, 36^, and metastasis^37^.

Moreover, fibrotic Fibronectin deposition correlates with resistance to anti-TNFα treatment in Crohn’s Disease patients^38^. Thus, our findings suggest that while ILC1 may have an unexpected anti-inflammatory role in the gut, their accumulation in inflamed tissues could exacerbate IBD-associated comorbidities, and be an indicator for poor treatment response. This unexpected contextualization of intestinal ILC1 was enabled by our reductionist, modular, and synthetic culture system, which could be further exploited to dissect dynamic interactions between other inaccessible cells and tissues, in both development and disease.

## Materials and methods

### Establishment of murine SIO cultures

Organoid cultures were established by isolating intact small intestine crypts from 6-8 week female CD45.1 C57BL/6 mice following established protocols4 and propagated in Matrigel (Corning) in basal media (DMEM/F12; 2mM Glutamax; 10mM HEPES; 1x Antibiotic-Antimycotic; 1x N2 supplement; 1x B27 supplement; all ThermoFisher, and 1mM Acetyl-L-cycteine, Sigma) supplemented with EGF (50ng/ml, R&D) and 50µl/ml of supernatant from both R-spondin (RSpo1-Fc) and Noggin cell lines, passaged every 4-5 days. RSpo1-Fc cell line was a kind gift from Professor Calvin Kuo and the Noggin cell line was a kind gift of Hubrecht Institute.

### Murine intestinal lymphocyte isolation

Lamina propria ILC1 were isolated from small intestines of litter matched female RORγt-GFP reporter mice following established protocols35. In short, excess fat and Peyer’s Patches were removed from the intestine, which was then opened longitudinally and rinsed thoroughly in ice-cold PBS. Small (1cm) sections were incubated in epithelial cell removal buffer for 2×15min (5mM EDTA and 10 mM HEPES in HBSS (GIBCO)), then tissue was cut into small pieces for extensive digestion of the extracellular matrix (collagenase (500µg/mL), dispase (0.5U/mL), DNAse1 (500µg/ml), 2%FBS in HBSS (GIBCO)). Samples were filtered through a 40µm strainer in 10%FCS-DMEM10, then lymphocytes were isolated using a 80%/40% isotonic percoll density gradient separation (centrifuged at 900G for 25min, no acceleration or deceleration). The interphase between 40% and 80% percoll was collected, filtered, and prepared for FACS isolation of ILC1 without further enrichment. Next, lymphocytes were rinsed with PBS, then stained with fixable LIVE/DEAD UV (ThermoFisher) in PBS for 15min in the dark at 4°C. The dye was quenched with sorting buffer, then the Fc-receptor was blocked (CD16/CD32, clone 93) for 10min at 4°C, followed by extracellular staining following standard flow cytometry protocols (1µl antibody/100µl sorting buffer/5million cells unless otherwise indicated). FACS antibodies were sourced from eBioscience (with the exception of CD45) and were as follows: CD3-Fluor450 (RB6-8C5), CD5-Fluor450 (53-7.3), CD19-Fluor450 (eBio1D3), Ly6G-Fluor450 (RB6-8C5), CD45-BV510 (30-F11, bioLegend) CD127-APC (A7R34), KLRG1-PerCP/eFluor710 (2F1), NKp46-PE/Cyanine7 (29A1.4), NK1.1-PE (PK136). Cells were rinsed, and sorted on a 70µm nozzle after calculation of compensation, and acquisition of Fluorescence Minus One controls for Lineage, CD127, and Nkp46. Gating strategies are outlined in Supplementary figures.

### Murine ILC-organoid co-cultures

Approximately 1500-2500 murine ILC1 were seeded with ∼100 mechanically disrupted SIO crypts per well, resuspended in 30μl ice-cold Matrigel, pipetted onto pre-heated tissue culture plates (Nunclon) and incubated at 37°C for 15-20min prior to addition of pre-warmed basal media supplemented with 50mM B2ME (R&D), 20ng/ml rhIL-2 (Sigma), 20ng/ml rmIL-7 (R&D), and 1ng/ml IL-15 (R&D), with media changes every 2-4 days.

### Human iPSC-derived intestinal organoids

The healthy KUTE-4 female skin fibroblast-derived human iPSC (hiPSC) line (available from the European Collection of Authenticated Cell Cultures (karyotyped, passage 24-36)) was cultured on plates coated with 40µl/ml vitronectin in PBS (StemCell Technologies). E8 (Gibco) media was changed daily, pockets of differentiation were actively removed, and round, pluripotent colonies were passaged with Versene (Gibco) every 4-6 days, when 60-70% confluent, or before circular colonies began merging.

KUTE-4 hiPSC were differentiated into human small intestine organoids (HIO) following established protocols^18^. In short, hiPSC were patterned toward definitive endoderm in RPMI with daily increasing B27 (0.2%, 1%, 2%) and 100ng/ml ActivinA (R&D) for 3.5 days, then patterned towards midgut in RPMI+2%B27 with 3μM CHIR99021 (Wnt agonist, TOCRIS) and 500ng/ml recombinant FGF4 (R&D) for 4days. At this point, CDX2 colonies were picked using a 200µl pipette tip, replated in 35µl Matrigel, then matured in basal media with hEGF 100ng/ml, R&D) rh-Rspondin (500ng/ml, R&D), rh-Noggin (100ng/ml, R&D), and 2ng/ml IL-2 supernatant for at least 35 days prior to establishing co-cultures with hILC1 or encapsulation in synthetic hydrogels for aILC1 characterization.

### Human lymphocytes isolation from patient biopsies

Studies in human tissues received ethical approval from the London Dulwich Research Ethics Committee (REC reference 15/LO/1998). Informed written consent was obtained in all cases. Inflammatory status of IBD patients was diagnosed by a consultant, and 15-20 colonic biopsies were procured by endoscopy. These were cultured on rat tail collagen I coated 9mm x 9mm x 1.5mm Cellfoam matrices (Cytomatrix PTY Ltd) in complete media (RPMI with 10% FBS) with antibiotics (penicillin, streptomycin, metronidazole, gentamicin and amphotericin) for 48h following established protocols^39, 40^. Colonic lamina propria mononuclear cells (cLPMCs) were then isolated from the supernatant ready for evaluation^41^ (protocol adapted from Di Marco Barros *et al*). Then, cLPMC were rinsed with PBS, treated with fixable Live/Dead-UV, and Fc blocked before being stained with CD45-eFluor450 (HI30; Invitrogen), Lineage cocktail 3-FITC (CD3, CD14, CD19, CD20; BD Biosciences), CD4-FITC (OKT4; BioLegend), TCRα/β-FITC (IP26; Biolegend), TCRγ/δ-FITC (B1; Biolegend), CD56-Alexa700 (B159; BD Pharmingen), CD7-PE-CF594 (M-T701; BD Horizon), CD127-PE-Cy7 (eBioRDR5; Invitrogen), c-kit-BV605 (104D2; BioLegend), CRTH2-PE (MACS Milltenyi Biotec), and CD161-APC (HP-3G10; BioLegend). cLPMC were sorted on a 70µM nozzle on Aria2 (BD). One biopsy per patient was fixed in 4%PFA and maintained for histology.

### Human ILC1-organoid co-cultures

Approximately 15-30 mature HIO were added to eppendorfs containing 50-300 hILC1 directly after FACS isolation from biopsies. The two components were centrifuged at 500G for 3min, supernatant was carefully removed, and the co-cultures were resuspended in 35µl Matrigel and plated onto pre-warmed tissue culture treated plates. The same culture conditions optimized for murine co-cultures were used for HIO-hILC1 co-cultures, including 50mM B2ME (R&D), 20ng/ml rhIL-2 (Sigma), 20ng/ml rmIL-7 (R&D), and 1ng/ml IL-15 (R&D), with media changes every 3-4 days.

### Cell isolation from co-cultures

After 4 days of murine and 7 days of human co-culture, Matrigel was disrupted and cells were collected into 15ml falcon tubes. For murine ILC1 cultures Matrigel disruption was not required, and cells were gently rinsed from the bottom of the plate using PBS+2%FCS. Samples were rinsed with PBS, then dissociated in TrypLe (Gibco) for 20mins at 37°C. The sorting buffer after this step contained DNAse (250µg/ml), EDTA (1μl/ml), and HEPES (1μl/ml) to maintain single epithelial cells and avoid clumping. Cells were titruated gently, centrifuged and resuspended in sorting buffer. Cells were then filtered (70µm), having pre-coated the filter with sorting buffer to minimize cell loss, and either rinsed with PBS for fixable Live/Dead staining (UV or nearInfraRed, Thermofisher), or stained with EpCAM, CD45, and the requisite combination of antibodies for the experiment, and analyzed (BD Fortessa) or sorted (BD ARIA3 Fusion & BD Aria 2). Isolation of murine IEC and ILC1 following co-culture was performed using EpCAM–APC Cy7 (G8.8, BioLegend), CD45-BV510 (30-F11, bioLegend), NK1.1 BV605 (PK136, BioLegend), CD44-PE (IM7, BioLegend). Isolation of human IEC, FB, and hILC1 used fixable Live/Dead-UV or Live/Dead-nIR CD45-eFluor450(HI30) Invitrogen, EpCAM-FITC (9C4; BioLegend), CD90-PE/Dazzle (Thy1; BioLegend)

### Flow cytometry

Flow cytometry data were acquired on a BD Fortessa2 and analyzed using FlowJo v10.5.3.

### RT-qPCR

For RNA isolation, cells were FACS-sorted directly into 250μl RLT (Qiagen) lysis buffer supplemented with 10μl/ml BME to stabilize the RNAse-rich intestinal epithelial tissue lysate. RNA was isolated using the RNeasy MicroRNA isolation kit (QIAGEN), and cDNA produced using RevertAid (Fisher), using oligo dTTTTT primers. Fast SYBR-green mix (Applied Biosystems) based RT-qPCR were run on a CFX384 TouchTM Real-Time PCR Detection System (BioRad), with no-template controls (NTC) and meltcurves for quality control, or using TaqMan Gene Expression Master Mix (Applied Biosystems) with FAM-probes, using an annealing temperature at 60°C for 39 cycles. All kits were used following manufacturers’ instructions. Primers were designed using PrimerBank and verified via BLAST against the mus musculus or homo sapiens genome on ensemble.org. Cq values were normalized to the housekeeping genes Hprt1 or GAPDH for SYBR and HPRT1 for TAQ probes.

#### Mouse SYBR

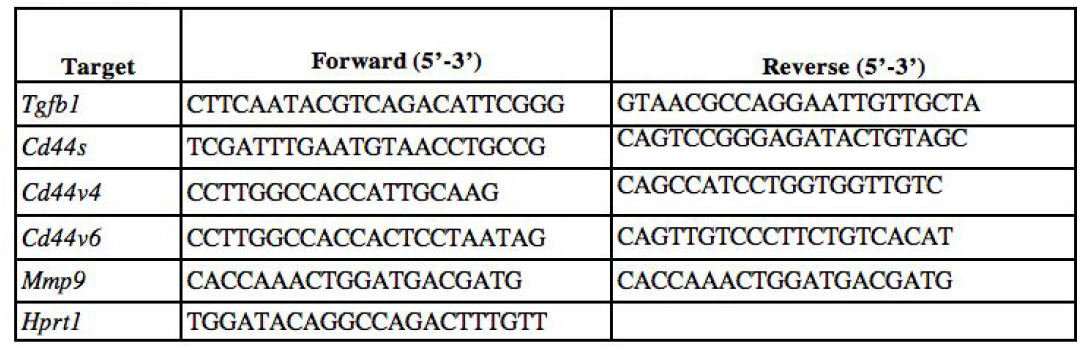

#### Human SYBR

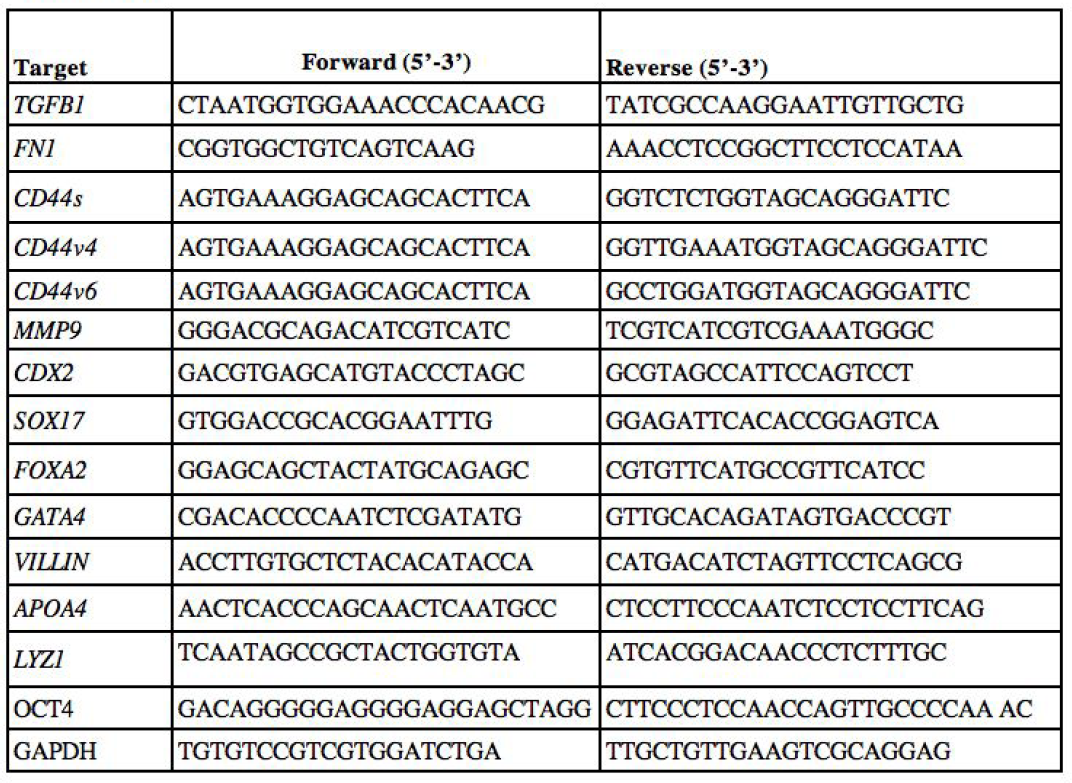

#### Human TAQ

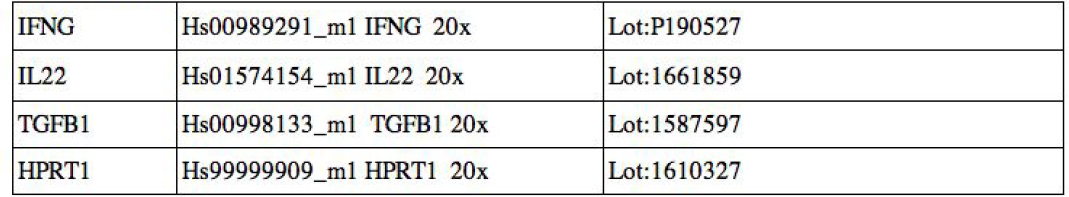

|  |  |  |
| --- | --- | --- |
| IFNG | Hs00989291_m1 IFNG 20x | Lot:P190527 |
| IL22 | Hs01574154_m1 IL22 20x | Lot:1661859 |
| TGFB1 | Hs00998133_m1 TGFB1 20x | Lot:1587597 |
| HPRT1 | Hs99999909_m1 HPRT1 20x | Lot:1610327 |

### Cytokine quantification

TGFβ-1 concentration in supernatant from 4day ILC1 co-cultures and SIO only controls was measured using the Mouse TGFβ-1 DuoSet ELISA (R&D Systems) with modified manufacturer’s instructions, whereby 100µl supernatant was incubated with the capture antibody overnight on a shaker at 4°C, not for 2h at RT. Optical density was measured in a plate reader (BioRAD) at 450nm, with correction at 540nm. Concentrations were obtained based on a regression equation (multimember regression, order 3) from the standard curve values (calculated in Excel).

Cytometric Bead Array for Th1/Th2/Th17 cytokines was obtained from BD biosciences and performed on 10µl supernatant after 4day co-culture following manufacturers’ instructions, using a BD Fortessa2, and analysed following manufacturers’ template based on a standard curve for each cytokine.

### PEG-peptide conjugate synthesis/characterization and hydrogel formation

Custom-designed peptides (supplied as either trifluoroacetic acid or acetate salts) used to either create A2+B4 (Ac-CREW-ERC-NH2) or A4+B4 designs containing either a degradable (Ac-GRDSGK-GPQG↓IWGQ-ERC-NH2), non-adhesive/non-degradable (non-adh/non-deg, Ac-KDW-ERC-NH2) or adhesive sequence (RGD, presented in either a linear Ac-RGDSGK-GDQGIAGF-ERC-NH2 or loop configuration (RGDSGD)K-GDQGIAGF-ERC-NH2) were synthesized by Peptide Protein Research, Ltd (UK) (all >98% purity). To create PEG-peptide conjugates, peptide was dissolved in anhydrous dimethyl sulfoxide (DMSO) at 10mg/ml and anhydrous triethylamine (TEA) (both Sigma) was added stoichiometrically to convert the peptide salts into their free forms in order to deprotonate the primary amine from the lysine side chain. Peptides were then conjugated to star-shaped, 4-arm PEG activated at each terminus with nitrophenyl carbonate (PEG-4NPC) by a nucleophilic substitution reaction between the primary amine on the side chain of the lysine residue of each peptide and NPC esters forming stable carbamate linkages. To accomplish this, a 16.67mg/ml solution of 10K PEG-4NPC (JenKem Technology, USA) in DMSO was reacted with peptide on an orbital shaker at either a 12:1 ratio of excess peptide to PEG-4NPC at RT for 30min (non-adh/non-deg), a 10:1 ratio at 60°C for 3h (degradable), a 8:1 ratio at RT for 2h (cyclic adhesive), or a 4:1 ratio at RT for 30min (linear adhesive). Conjugates were then snap frozen on dry ice and lyophilized. To reduce disulfide bonds, conjugates were dissolved in carbonate-bicarbonate buffer at pH9.0 and treated with DTT (0.1g/ml) for 6h at RT, after nitrogen purging (molar ratio of 4.5:1, DTT:peptide). Conjugates were then purified 4x in MiliQ water using Merck Millipore Ultrafiltration 1MWCO units (10KDa cut-off), snap frozen and lyophilized again prior to storage at −20°C

Conjugation conversion was determined for non-adh/non-deg, degradable and cyclic adhesive conjugates by size exclusion chromatography (SEC) using a Gilson HPLC system. Calibration was performed using standards of known peptide concentration. The relative amount of unreacted peptide was assessed by estimating the concentration of free peptide in the crude reaction mixture. We observed that ∼80-100% of PEG arms were conjugated with peptide (conjugation efficiency).

Hydrogels with an A4+B4 design were formed by reacting PEG-peptide conjugates with star-shaped 4-arm PEG (20kDa, unless otherwise noted) bearing vinyl sulfone groups at each chain terminus (PEG-4VS). The reaction was performed in a stoichiometric ratio of 1:1 in 30 mM HEPES buffer (pH8.0, with 1X HBSS in a desired volume) through a Michael-type reaction between a cysteine thiol on the C-terminal of the peptide with the vinyl-sulfone group on PEG-4VS. To form 2.5% non-adh/non-deg hydrogels for swelling and rheological studies using an A2+B4 design, Ac-CREW-ERC-NH2 was reacted with PEG-4VS in a stoichiometric ratio of 2:1 in 30mM HEPES buffer (pH8.0). Hydrogels were then allowed to form for 45-60min prior to being placed in PBS/culture media as indicated.

The conjugation efficiency of the thiol vinyl sulfone reaction was determined using proton NMR and Ellman’s assay. For 1H NMR experiments, PEG-peptide conjugate and PEG-4VS were dissolved separately in HEPES buffer and lyophilized. The resulting powders were dissolved separately in deuterium oxide to a final polymer/peptide concentration of 1.5 wt%, mixed at stoichiometric ratio and loaded into 0.3mm diameter NMR tubes. Acquisition of spectra was performed on a Bruker 700MHz NMR spectrometer. The first measurement was made after 6 min and additional measurements collected for up to 1h when aromatic signals from the vinyl sulfone were no longer distinguishable. Hydrogel formation was observed inside the NMR tubes at the end of the experiment.

The relative quantity of free thiols during the reaction were quantified using the molar absorptivity of Ellman’s reagent, as previously described. Briefly, a 4 mg/ml solution of Ellman’s reagent was prepared in reaction buffer (0.1M PBS, pH8.0, containing 1mM EDTA). Hydrogels were prepared and the cross-linking reaction halted after 5, 10, 15, 30 and 60min using a 1:50 dilution of reaction buffer and Ellman’s reagent solution. Samples were then incubated for 15min and absorbance measured at 412nm. Free thiols in a peptide (Ac-KDWERC-NH2) solution alone were quantified using the same method. The concentration of free thiols was calculated based on the molar extinction coefficient of Ellman’s reagent (14150 M-1 cm-1) and Equation (Eq. 1)

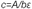

Where A is the absorbance of the sample at 412nm, b is 1cm and ε is the molar extinction coefficient.

### Molecular dynamics simulations

Coarse-grain classical molecular dynamics simulations were used to study the cross-linking of hydrogels formed with either A4+B4 or A2+B4 designs. Three hydrogel systems were simulated in replicate: Ac-KDWERC-NH2 and H-SREWERC-NH2 are A4+B4 designs and Ac-CREWERC-NH2 is a A2+B4 design. As in our experimental work, Ac-KDWERC-NH2 and H-SREWERC-NH2 are ‘pre-conjugated’ in the simulation to PEG-4NPC and the reaction with PEG-4VS is simulated. For Ac-CREWERC-NH2, there was no ‘pre-conjugation’ step and PEG-4VS was allowed to react with the free peptide. Ac-KDWERC-NH2 is the non-adh/non-deg peptide used in experimental studies; however, as the net charge of Ac-KDWERC-NH2 is −1, while that of Ac-CREWERC-NH2 is 0, we also simulated the A4+B4 system with H-SREWERC-NH2 (net charge of 0) to study our system independently of electrostatic bias in bond formation. Supplementary Fig. 14 summarizes the molecules and water beads used in each simulated system. As we have used the MARTINI forcefield to represent the peptide cross-linkers, ions and the water molecules, each water bead represents 4 water molecules. The PEG molecules were modelled with the MARTINI-like forcefield, as described in Lee et al. (2009)^39, 42^.

The initial systems were built using PACKMOL^40^ to randomly place each component within a 40nm × 40nm × 40nm simulation box. The LAMMPS simulation engine was used for all simulations^43^. The software package Moltemplate (<https://moltemplate.org/>), was used to^43^ convert the configurations generated by PACKMOL to those readable by LAMMPS. The resulting systems have PEG concentrations of 2.5%. Once the initial systems were built, we first minimized energy using the steepest descent algorithm with an energy tolerance of 1 × 10-4 and a force tolerance of 1 × 10-6. The systems were then equilibrated by carrying out a series of simulations with the NVT (constant number of particles, volume and temperature) ensemble with the Langevin thermostat and a target temperature of 300K. During these simulations, the systems were run for 1ps with a 1fs timestep, 3ps with a 3fs timestep, 10ps with a 10fs timestep and then 400ps with a 20fs timestep. The volume was then equilibrated by carrying out a series of simulations with the NPT (constant number of particles, pressure and temperature) ensemble employing the Langevin thermostat and the Parrinello-Rahman barostat. In these simulations, the time step was again increased (1ps with a 1fs timestep, 3ps with a 3fs timestep, 10ps with a 10fs timestep, and 2ns with a 20 fs timestep). The densities of the systems were then equilibrated using the NPT ensemble with a Nosé-Hoover thermostat and barostat for a simulation lasting 2ns with a 20fs timestep. Finally, we equilibrated the temperature of the simulated systems to 450K using an NVT simulation with the Nosé-Hoover thermostat which lasted 40ns with a 20fs timestep.

Production simulations were carried out in the NVT ensemble at 450K (to increase diffusion and allow for investigation of hydrogel cross-linking in a reasonable amount of simulation time) with a timestep of 20fs. During simulations, we employed cross-linking methods that have been previously shown in simulations to lead to the formation of hydrogels^44^. In short, we identify beads that can react with one another and then check at regular intervals of time (treact) if any two reaction partner beads are within a given distance (rreact) of one another. If so, then a new bond is formed with a given probability (preact). Here, we used a bead containing the sulphur atom within the cysteine residue on each peptide as a reactive bead in the simulations. Its reaction partner was the terminal bead on each arm of PEG-4VS. This reaction model is consistent with the chemistry that forms the hydrogels. The reactions were modelled using treact = 20ps, rreact = 5.0Å and preact = 0.5. Once a bond was formed between any cysteine bead and a terminal bead on a PEG-4VS, neither of those beads could form any other new bonds during the simulation. Production simulations were run for at least 5.5µs seconds for 1 replica of each system, and the other replica was run for ∼4µs.

### Measurements of hydrogel swelling

30uL hydrogels were formed in Sigmacote®-treated 6mm diameter glass cylindrical moulds and submersed in PBS. Hydrogel weight was monitored, and the wet weight measured once swelling equilibrium had been achieved (after 48h). Hydrogels were then lyophilized to determine dry weight and the mass swelling ratio (Qm) calculated using the following equation:

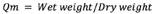

### Rheological measurements of hydrogel gelation

Hydrogel gelation was assessed on a strain-controlled ARES from TA Instruments using a 25mm cone with a 0.02rad cone-and-plate by carrying out small amplitude oscillatory time sweep measurements at a strain of 5% and a constant angular frequency of 1rad/s. All measurements were carried out at 37°C, sealing the chamber with oil to prevent evaporation. To perform measurements, 80µL hydrogels were placed in the instrument and storage modulus G’ and loss modulus G’’ were recorded as a function of time in TRIOS software. Subsequently an amplitude sweep was carried out, recording G’ and G” as a function of shear amplitude in the range of 1-100% shear strain, to determine the linear viscoelastic region. Finally, a frequency sweep was recorded, measuring G’ and G” as a function of shear frequency in a range of 100 - 0.1rad/s, to assess the hydrogels’ temporal behaviour.

### Quantification of MMP-mediated hydrogel degradation

Degradability was assessed on 30µL hydrogels formed with either 0, 45, 75, or 100% of PEG-peptide conjugates containing a degradable sequence (all other cross-links formed with non-adh/non-deg peptides) that had been allowed to swell in PBS for 24h. To degrade hydrogels, PBS was replaced with a solution of TCNB buffer (50mM Tris, pH7.5, with 100mM NaCl, 10mM CaCl2) containing 89.5 nM human MMP9 (Sigma SAE0078) and incubated at 37°C. Degradation was determined by measuring the absorbance of tryptophan found on the cleaved peptide section in the supernatant at 280nm. Degradation was determined by calculating the ratio of the cleaved peptide in solution to that in the initial hydrogel.

Mechanical testing by atomic force microscopy (AFM)-based force spectroscopy 30µL hydrogels were formed in Sigmacote®-treated 6mm diameter glass cylindrical moulds in 35mm petri dishes and stored in PBS at 4°C prior to testing. Force-distance measurements were carried out on a JPK Nanowizard 4 (JPK instruments AG, DE) directly on hydrogels immersed in PBS at RT. To perform indentation measurements, spherical glass beads (diameter 10μm; Whitehouse Scientific, UK) were mounted onto tipless triangular silicon nitride cantilevers (spring constant (K) ≈ 0.12N m−1; Bruker AXS SAS, FR) using UV-cross-linked Loctite super glue. The deflection sensitivity of the AFM photodiode was then calibrated by collecting a single force-distance curve on a glass slide. Cantilevers were calibrated using the thermal method^45^ in air. Measurements were made on 6 different locations across each hydrogel’s surface (100µm x 100µm areas, 100 force curves per location on 3 independent hydrogels per condition). Indentations were carried out with a relative setpoint force of 3nN and a loading rate of 4μm s−1. Data were collected using JPK proprietary software (JPK Instruments AG, DE). The Oliver–Pharr model for a spherical tip was used to determine E. Outliers were removed using a ROUT test (Q=1%). As for other hydrated biological samples, we assumed that volume was conserved and assigned a Poisson’s ratio of 0.5.

| Polymer Concentration |  |  |  |  |
| --- | --- | --- | --- | --- |
| 10% | 5% | 3% | 2.5% | 2 % |
| 1640 | 1609 | 1781 | 1613 | 1506 |

Table: Total number of measurements (force curves) collected for AFM-based measurements of stiffness. Measurements were made on 3 independent hydrogels per condition.

### Human ancillary ILC1 hydrogel co-cultures

HIO were harvested, titruated, and thoroughly rinsed with ice cold PBS, then resuspended in pH8.0 buffered phenol free and protein free HBSS (pH8-HBSS, Gibco), spun down, resuspended and left on ice until encapsulation. PEG-peptide conjugates were dissolved in ice cold pH8-HBSS, vortexed and centrifuged, and combined based on the gel composition (e.g. IM-DEG: 20% cyclic adhesive, 45% DEG, 35% NON-DEG). An equal molar mass of PEG-4VS was weighed in a protein-low-binding eppendorf, dissolved in pH8-HBSS, centrifuged and added to the PEG-conjugate mix, and the sample was vortexed and centrifuged again. HIO were then rapidly mixed into the PEG-peptide conjugate/PEG-4VS mix using protein-low binding 200μl tips, and pipetted into a pre-warmed, Sigmacote®-treated glass ring in a 24-well plate Nunclon well. HIO-laden gels were incubated at 37°C and after 30min the glass ring was removed using an autoclaved forceps, and basal media supplemented with 50mM BME, 20ng/ml IL-2, 20ng/ml IL-7, and 1ng/ml IL-15 (R&D) was added to the cultures.

### Stiffness mapping of IM-DEG hydrogel-HIO cultures by AFM

Organoids were encapsulated in IM-DEG hydrogels formed in Sigmacote®-treated 10mm diameter glass cylindrical moulds and submersed in culture media. Force-distance curves were collected on a JPK Nanowizard-CellHesion (JPK instruments AG, DE) mounted onto an inverted light microscope. Tipless triangular silicon nitride cantilevers (spring constant (K) ≈ 0.12 N m−1; Bruker AXS SAS, FR) were calibrated using the thermal method^45^ in air and then functionalized with 50μm silica beads (Cospheric, USA) as above. Prior to measurements, the deflection sensitivity of the AFM photodiodes was calibrated by collecting a single force-distance curve on a glass slide in liquid.

Prior to measurements, cultures containing organoids were placed in CO2 independent media (Sigma). Maps were then collected across the surface of hydrogels in regions where an organoid could be clearly identified on the inverted light microscope. Map sizes varied depending on the organoid size but indentations were done in either 8×8 or 16×16 grids with the largest map being 300×300µm and the smallest 150×150µm. For each map, indentations were carried out at a relative setpoint of 2.5nN and a loading rate of 4μm s−1. The manufacturer’s proprietary JPK SPM software 6.1 (JPK Instruments AG, DE) was used to determine E using the Hertz model for a spherical tip. As for other hydrated biological samples, we assumed that volume was conserved and assigned a Poisson’s of 0.5.

### Multiple particle tracking microrheology

0.75 μm diameter fluorescent beads (Fluoresbrite YG Carboxylate Kit I 21636-1, Polysciences Inc.) were suspended at a concentration of 0.04% (w/w) in the polymer solution within 5-10µl HIO-laden hydrogels with or without aILC1. Samples were prepared and imaged in Ibidi slide-chambers (μ-Slide Angiogenesis, 81501) using a modified setup and analysis pipeline to that previously described by Schultz et al.^28^. Approximately 7-10 HIO were embedded in each hydrogel.

Time-lapses of moving beads were acquired using an Olympus TIRF System using an excitation of 488nm. Time-lapses of 800 frames were collected at a rate of 16.9ms per frame and exposure time of 1.015ms. HIO images were also captured in brightfield. ImageJ TrackMate was used to segment beads and create trajectories across the 800 frames. Mean squared displacements of individual beads were then calculated from the TrackMate output using custom R-script (Supplementary Data Set 2).

### Immunocytochemistry

Co-cultures were fixed for 10min using 4%PFA and either stained as whole organoids in FCS-coated Eppendorf tubes, or cryoprotected (overnight 30% glucose for organoids in Matrigel, overnight OCT replacement for hydrogels) and embedded in OCT for cryosectioning on a Penguin cryostat. Images were acquired on an inverted Leica SP8 inverted confocal microscope. Cells were blocked with 2%FCS and 0.05%TRITON-X in PBS for 1h at RT, stained at 4°C overnight, and in secondaries (ThermoFisher) for 1h at RT, followed by extensive rinse steps. Heat Antigen Retrieval was performed in pH8.0 basic conditions (10min, 95°C waterbath) for CD44v6 staining to reveal the v6 epitope, and optimize signal strength. All secondary antibodies were AlexaFluor conjugate dyes (488, 555, or 647) raised in donkey (ThermoFisher) Image processing and quantification was performed using FIJI (ImageJ). Image quantification was performed on max-intensity projections with the same (i) number of Z-stacks and (ii) the same brightness and contrast settings in each fluorophore channel, having been taken with the same laser power and gain values. Background intensity of the channel was subtracted from average intensity, which was then normalized to DAPI (nuclear) intensity. Nuclear phosphorylated p38 (p-p38) was quantified as follows: DAPI channel was processed to “binary” and erosion (E) and dilation (D) operations were performed to homogenize the nucleus area (E, D, D, E). An overlay with the outlines of the nuclei was created and saved in the ROI manager, which was then superimposed to either p-p38 channel. The mean intensity of each fluorophore within the defined nuclei areas was measured, giving an approximate measure of intensity values/nuclei.

Antibodies used were as follows:

| Target | Host species | Provider | Cat# | Reactivity | Dilution |
| --- | --- | --- | --- | --- | --- |
| Fibronectin | Rabbit | abcam | Ab2413 | Human, Mouse | 1:1000 |
| CD45 (HI30) | Mouse | BioLegend | 304001 | Human | 1:500 |
| FoxA2 | Rabbit | Abcam | Ab108422 | Human | 1:100 |
| CDX2 | Rabbit | Abcam | Ab76541 | Human | 1:250 |
| Lysozyme | Rabbit | Invitrogen | PA5-16668 | Human, Mouse | 1:500 |
| E-Cadherin<br>(DECMA-1) | Rat, (Alexa647) | eBioscience | 51-3249-82 | Human | 1:250 |
| β-catenin | Mouse | BD<br>Biosciences | 610153 | Human, Mouse | 1:100 |
| CD44 (IM7) | Rat | eBioscience | 14-0551-82 | Human, Mouse | 1:250 |
| EpCAM | Rabbit | abcam | Ab71916 | Human, Mouse | 1:750 |
| GATA4 | Mouse | Santa Cruz | Sc-25310 | Human | 1:500 |
| SOX17 | Goat | R&D | Af1924 | Human | 1:500 |
| Vimentin | Chicken | Millipore | Ab5733 | Human, Mouse | 1:1000 |
| TGFβR1 | Rabbit | Santa Cruz | Sc-339 | Human, Mouse | 1:500 |
| ZO-1 | Rabbit | ThermoFisher | 61-7300 | Human, Mouse | 1:100 |
| Phos- p38 (Thr180/Tyr182) | Rabbit | Cell Signalling | 4511 | Human, Mouse | 1:250 |
| Phalloidin-TexRed<br>(F-Actin) | N/A | ThermoFisher | T7471 | Human, Mouse | 1:5000 |
| DAPI | N/A | ThermoFisher | D1306 | Human, Mouse | 1:5000 |
| CD44var6 (VFF-18) | mouse | ThermoFisher | BMS125 | Human | 1:500 |
| CD44v6 (9A4) | Rat IgG1 | BioRad | MCA1967 | Mouse | 1:500 |
| SMA | Mouse IgG<sub>2A</sub> | R&D | MAB142 | Human/Mouse/Rat | 1:1000 |
| beta-catenin | goat | R&D | AF1329 | Human/Mouse/Rat | 1:500 |

### Viability tests

Viability staining were either performed by measuring the % of DAPI negative cells using a flow cytometer, or were performed using 5mg/ml Fluorescein Diacetat (FDA; Sigma-Aldrich Co. LLC, C-7521) and 2mg/ml Propridium Iodide (Sigma-Aldrich Co. LLC, P4170) for 90 seconds, followed by two rinse steps with PBS and acquisition on a Leica SP8 confocal microscope.

### Production of RNAseq dataset

The cells were harvested as described above for sorted by flow cytometry (BD ARIA3 Fusion) into RLT (Qiagen) lysis buffer. RNA was harvested using RNeasy MicroRNA isolation kit (QIAGEN), and RIN values were assessed using RNA 6000 Pico Kit (Agilent). The library was prepared using SMARTSeq2 and sequenced by Illumina HiSeq 4000 at the Wellcome Trust Oxford Genomics Core, where basic alignment (GRCm38.ERCC (2011)) and QC was also performed.

### RNAsequencing data analysis

Exploratory data analysis and filtering: The data count matrix was filtered for genes with a mean of less than 3 to remove very low count genes, and genes where most of the counts were zero were also removed. Model Description: A varying intercepts hierarchical modelling framework was used to model the expression for each gene. This analysis was implemented in R^46^ and Stan^47^. Pathway Analysis: Gene set enrichment analysis (GSEA) was carried out using the R package GAGE, and predicted upstream regulators, canonical pathways, and diseases and functions were determined using IPA (Qiagen) of padj<0.05 genes, excluding chemicals.

### hMSC attachment on 2D hydrogel surfaces

Human bone marrow-derived stromal cells (hMSC) were obtained from the Imperial College Healthcare Tissue Bank (ICHTB, HTA license 12275). ICHTB is supported by the National Institute for Health Research (NIHR) Biomedical Research Centre based at Imperial College Healthcare NHS Trust and Imperial College London. ICHTB is approved by the UK National Research Ethics Service to release human material for research (12/WA/0196) as previously described^48^. The samples for this project were issued from sub-collection R16052. 50 µL 5% hydrogels formed with 5K PEG-4VS were formed in 6-well plates and 24mm Sigmacote®-treated coverslips placed on top. After hydrogel formation, 5,000 hMSC/cm2 were seeded and allowed to adhere for 2h prior to the addition of basal culture media. After 24h, hMSC were fixed in 4%PFA, permeabilized in 0.2% (v/v) Triton X-100 and stained with Phalloidin-TRITC (Sigma) and DAPI. Cells were imaged on an Olympus inverted fluorescent microscope equipped with a Jenoptik Camera.

### Animals

CD45.1 mice (B6.SJL-PtprcaPepcb/BoyCrl, Charles River) and Rorc(γt)-GfpTG reporter mice54 (a kind gift from Gerard Eberl) were housed under specific-pathogen-free conditions at accredited Charles River and King’s College London animal units in accordance with the UK Animals (Scientific Procedures) Act 1986 (UK Home Office Project License (PPL:70/7869 to September 2018; P9720273E from September 2018).

### Statistics

Statistical analyses were performed in GraphPad Prism version 8.1.2.

## Acknowledgements

G.M.J. acknowledges a Ph.D. fellowship from the Wellcome Trust (203757/Z/16/A) and a BRC Bright Sparks Precision Medicine Early Career Research Award. E.G. acknowledges a Philip Leverhulme Prize from the Leverhulme Trust. J.F.N. acknowledges a Marie Skłodowska-Curie Fellowship, a King’s Prize fellowship, a RCUK/UKRI Rutherford Fund fellowship (MR/R024812/1) and a Seed Award in Science from the Wellcome Trust (204394/Z/16/Z). M.D.A.N. is supported by a PhD studentship funded by the BBSRC London Interdisciplinary Doctoral Programme. E.R. acknowledges a Ph.D. fellowship from the Wellcome Trust (215027/Z/18/Z). S.L. gratefully acknowledges the UK Medical Research Council (MR/N013700/1) for funding through the MRC Doctoral Training Partnership in Biomedical Sciences at King’s College London. G.M.L. is supported by grants awarded by the Wellcome Trust (091009) and the Medical Research Council (MR/M003493/1 & MR/K002996/1). N.J.W. acknowledges a Jane & Aatos Erkko Foundation Personal Scholarship. R.M.P.dS acknowledges a King’s Prize fellowship supported by the Wellcome Trust (Institutional Strategic Support Fund), King’s College London and the London Law Trust. Via C.D.L.’s membership in the UK’s HEC Materials Chemistry Consortium, which is funded by EPSRC (EP/L000202, EP/R029431), this work used the ARCHER UK National Supercomputing Service (http://www.archer.ac.uk) and the UK Materials and Molecular Modelling Hub (MMM Hub) for computational resources, which is partially funded by EPSRC (EP/P020194/1), to carry out the MD simulations. The authors also wish to thank the BRC flow cytometry core team, and acknowledge financial support from the Department of Health via the NIHR comprehensive Biomedical Research Centre award to Guy’s and St. Thomas’ NHS Foundation Trust in partnership with King’s College London and King’s College Hospital NHS Foundation Trust. The views expressed are those of the author and not necessarily those of the NHS, the NIHR, or the Department of Health. The authors thank Camilla Dondi, Daniel Foyt, and Oksana Birch for technical assistance. The authors are grateful to Dr. Jo Spencer and Dr. Kelly Schultz for helpful conversations about CD44 and microrheology, for technical support from Drs. Rebecca Beavil and Andrew Beavil with SEC-HPLC, Dr Hugo Sinclair from the Microscopy Innovation Centre for assistance acquiring micro-rheology data, Dr Richard Thorogate and the London Centre for Nanotechnology for assistance with AFM, Dr R. A. Atkinson and the NMR Facility of the Centre for Biomolecular Spectroscopy at King’s College London, which was established with awards from the Wellcome Trust, British Heart Foundation and King’s College London for assistance with NMR, and Simon Engledow at the Oxford Wellcome Genomics Centre for processing the RNAseq samples. Finally, we would like to thank Dr. Luke Roberts, Erin Slatery, and Dr. Rocio Sancho for critically reading this manuscript and providing helpful feedback.

## Author contributions

G.M.J., M.D.A.N., T.T.L.Y., L.B., J.F.N. and E.G. developed experimental protocols, conducted experiments, and analyzed data. T.T.Y.L., J.H., O.P.O., N.J.W., C.A.D., N.D.E., RMP.dS. designed and synthesized the hydrogel. R.M.P.dS. designed the peptide sequences. M.D.A.N., D.H., S.L., T.T.L.Y., G.M.J. and D.M., characterized the hydrogel. S.L. and C.D.L. performed the molecular dynamics simulations. G.M.J. and J.F.N. designed the RNA-sequencing experiment, and U.N. and M.C. provided bioinformatic analysis. G.M.J., P.R., E.R., and T.Z. performed lymphocyte and SIO isolations. G.M.J. conducted and analysed murine and human experiments in Matrigel. M.D.A.N., G.M.J., E.H. designed and conducted microrheology and AFM experiments. G.M.L., O.O., and D.D. contributed reagents, biopsies, and hiPSC cell lines. G.M.J., E.G. and J.F.N. conceived the ideas, initiated the project, interpreted the data, and prepared the manuscript. E.G. and J.F.N. supervised the project. All authors revised the manuscript.

## Data availability

The differentially expressed genes identified in the RNAsequencing dataset are available in Supplementary Data Set 1. The data have also been deposited with GEO, at https://github.com/uhkniazi/BRC_Organoids_Geraldine, and Research Data (Springer Nature) with the dataset identifiers made available upon publication. All other data supporting the findings of this study are available within the article and its supplementary information files or from the corresponding author upon reasonable request.

## Code availability statement

All code used to analyse the molecular dynamics simulations were tools that were built in-house. All codes with accompanying documentation as to how to use them will be made available at https://github.com/Lorenz-Lab-KCL and https://nms.kcl.ac.uk/lorenz.lab/wp/ upon publication and will be freely accessible. R code for determining alpha from MSD data for microrheology is available in Supplementary Data Set 2 with its doi made available upon publication.

## References

1. Haber, A. L. et al. A single-cell survey of the small intestinal epithelium. Nature 551, 333–339 (2017).

2. Vivier, E. et al. Innate Lymphoid Cells: 10 Years On. Cell 174, 1054–1066 (2018).

3. Lindemans, C. et al. Interleukin-22 promotes intestinal-stem-cell-mediated epithelial regeneration. Nature 528, 560 (2015).

4. Spits, H., Bernink, J. H. & Lanier, L. NK cells and type 1 innate lymphoid cells: partners in host defense. Nature Immunology 17, 758–764 (2016).

5. Bernink, J. H. et al. Human type 1 innate lymphoid cells accumulate in inflamed mucosal tissues. Nature Immunology 14, 221–229 (2013).

6. Pagnini, C., Pizarro, T. T. & Cominelli, F. Novel Pharmacological Therapy in Inflammatory Bowel Diseases: Beyond Anti-Tumor Necrosis Factor. Front. Pharmacol. 10 (2019).

7. Sato, T. et al. Paneth cells constitute the niche for Lgr5 stem cells in intestinal crypts. Nature 469, 415–418 (2011).

8. Spence, J. R. et al. Directed differentiation of human pluripotent stem cells into intestinal tissue in vitro. Nature 470, 105–9 (2011).

9. Senbanjo, L. T. & Chellaiah, M. A. CD44: A Multifunctional Cell Surface Adhesion Receptor Is a Regulator of Progression and Metastasis of Cancer Cells. Frontiers in cell and developmental biology 5, 18 (2017).

10. Martin-Gallausiaux, C. et al. Butyrate produced by gut commensal bacteria activates TGF-beta1 expression through the transcription factor SP1 in human intestinal epithelial cells. Scientific Reports 8, 1–13 (2018).

11. Zeilstra, J. et al. Stem cell CD44v isoforms promote intestinal cancer formation in Apc(min) mice downstream of Wnt signaling. Oncogene 33, 665–670 (2014).

12. Ghatak, S. et al. Transforming growth factor β1 (TGFβ1) regulates CD44V6 expression and activity through extracellular signal-regulated kinase (ERK)-induced EGR1 in pulmonary fibrogenic fibroblasts. Journal of Biological Chemistry 292, 10465–10489 (2017).

13. Schmitt, M., Metzger, M., Gradl, D., Davidson, G. & Orian-Rousseau, V. CD44 functions in Wnt signaling by regulating LRP6 localization and activation. Cell death and differentiation 22, 677–689 (2015).

14. Ribes, B. M. et al. Effectiveness and safety of pirfenidone for idiopathic pulmonary fibrosis. Eur J Hosp Pharm (2019).

15. Yin, N. et al. p38γ MAPK is required for inflammation-associated colon tumorigenesis. Oncogene 35, 1039–1048 (2016).

16. Fujii, M. et al. Human Intestinal Organoids Maintain Self-Renewal Capacity and Cellular Diversity in Niche-Inspired Culture Condition. Cell Stem Cell 23, 787–793.e6 (2018).

17. Dotti, I. et al. Alterations in the epithelial stem cell compartment could contribute to permanent changes in the mucosa of patients with ulcerative colitis. Gut 66, 2069–2079 (2017).

18. McCracken, K. W., Howell, J. C., Wells, J. M. & Spence, J. R. Generating human intestinal tissue from pluripotent stem cells in vitro. Nature Protocols 6, 1920–1928 (2011).

19. Jung, K. B. et al. Interleukin-2 induces the in vitro maturation of human pluripotent stem cell-derived intestinal organoids. Nature Communications 9, 1–13 (2018).

20. Shimshoni, E., Yablecovitch, D., Baram, L., Dotan, I. & Sagi, I. ECM remodelling in IBD: innocent bystander or partner in crime? The emerging role of extracellular molecular events in sustaining intestinal inflammation. Gut 64, 367–372 (2015).

21. Farkas, K. et al. The Diagnostic Value of a New Fecal Marker, Matrix Metalloprotease-9, in Different Types of Inflammatory Bowel Diseases. Journal of Crohn’s and Colitis 9, 231–237 (2015).

22. Hughes, C. S., Postovit, L. M. & Lajoie, G. A. Matrigel: A complex protein mixture required for optimal growth of cell culture. PROTEOMICS 10, 1886–1890 (2010).

23. Ehrbar, M. et al. Biomolecular Hydrogels Formed and Degraded via Site-Specific Enzymatic Reactions. Biomacromolecules 8, 3000–3007 (2007).

24. Zhong, M., Wang, R., Kawamoto, K., Olsen, B. D. & Johnson, J. A. Quantifying the impact of molecular defects on polymer network elasticity. Science (New York, N.Y.) 353, 1264–8 (2016).

25. Cruz-Acuña, R. et al. PEG-4MAL hydrogels for human organoid generation, culture, and in vivo delivery. Nature Protocols 13, 2102–2119 (2018).

26. Gu, Y. et al. Semibatch monomer addition as a general method to tune and enhance the mechanics of polymer networks via loop-defect control. Proceedings of the National Academy of Sciences of the United States of America 114, 4875–4880 (2017).

27. Stewart, D. C. et al. Quantitative assessment of intestinal stiffness and associations with fibrosis in human inflammatory bowel disease. PLOS ONE 13, e0200377 (2018).

28. Schultz, K. M., Kyburz, K. A. & Anseth, K. S. Measuring dynamic cell–material interactions and remodeling during 3D human mesenchymal stem cell migration in hydrogels. PNAS 112, E3757–E3764 (2015).

29. Hocevar, B. A., Brown, T. L. & Howe, P. H. TGF-beta induces fibronectin synthesis through a c-Jun N-terminal kinase-dependent, Smad4-independent pathway. EMBO J. 18, 1345–1356 (1999).

30. Li, J., Wu, J., Zhang, M. & Zheng, Y. Dynamic changes of innate lymphoid cells in acute ST-segment elevation myocardial infarction and its association with clinical outcomes. Scientific Reports 10, 1–12 (2020).

31. Bauché, D. & Marie, J. C. Transforming growth factor β: a master regulator of the gut microbiota and immune cell interactions. Clinical & Translational Immunology 6, e136 (2017).

32. Fenton, T. M. et al. Inflammatory cues enhance TGFβ activation by distinct subsets of human intestinal dendritic cells via integrin αvβ8. Mucosal immunology 10, 624–634 (2017).

33. Bal, S. M., Golebski, K. & Spits, H. Plasticity of innate lymphoid cell subsets. Nature Reviews Immunology, 1–14 (2020).

34. Wang, J. et al. CD44v6 promotes β-catenin and TGF-β expression, inducing aggression in ovarian cancer cells. Mol Med Rep 11, 3505–3510 (2015).

35. Wang, Z., Zhao, K., Hackert, T. & Zöller, M. CD44/CD44v6 a Reliable Companion in Cancer-Initiating Cell Maintenance and Tumor Progression. Frontiers in Cell and Developmental Biology 6, 97 (2018).

36. Ma, L., Dong, L. & Chang, P. CD44v6 engages in colorectal cancer progression. Cell Death & Disease 10, 30 (2019).

37. Todaro, M. et al. CD44v6 Is a Marker of Constitutive and Reprogrammed Cancer Stem Cells Driving Colon Cancer Metastasis. Cell Stem Cell 14, 342–356 (2014).

38. de Bruyn, J. R. et al. Intestinal fibrosis is associated with lack of response to Infliximab therapy in Crohn’s disease. PLoS One 13 (2018).

39. de Jong, D. H. et al. Improved Parameters for the Martini Coarse-Grained Protein Force Field. Journal of Chemical Theory and Computation 9, 687–697 (2013).

40. Martínez, L., Andrade, R., Birgin, E. G. & Martínez, J. M. PACKMOL: A package for building initial configurations for molecular dynamics simulations. Journal of Computational Chemistry 30, 2157–2164 (2009).

41. Di Marco Barros, R. et al. Epithelia Use Butyrophilin-like Molecules to Shape Organ-Specific γδ T Cell Compartments. Cell 167, 203–218.e17 (2016).

42. Lee, H., de Vries, A. H., Marrink, S. & Pastor, R. W. A Coarse-Grained Model for Polyethylene Oxide and Polyethylene Glycol: Conformation and Hydrodynamics. The Journal of Physical Chemistry B 113, 13186–13194 (2009).

43. Plimpton, S. Fast Parallel Algorithms for Short-Range Molecular Dynamics. Journal of Computational Physics 117, 1–19 (1995).

44. Bode, F. et al. Hybrid gelation processes in enzymatically gelled gelatin: impact on nanostructure, macroscopic properties and cellular response. Soft Matter 9, 6986–6999 (2013).

45. Hutter, J. L. & Bechhoefer, J. Calibration of atomic-force microscope tips. Review of Scientific Instruments 64, 1868–1873 (1993).

46. Luo, W., Friedman, M. S., Shedden, K., Hankenson, K. D. & Woolf, P. J. GAGE: generally applicable gene set enrichment for pathway analysis. BMC Bioinformatics 10, 161 (2009).

47. Carpenter, B. et al. Stan: A Probabilistic Programming Language. Journal of Statistical Software 76, 1–32 (2017).

48. Ferreira, S. A. et al. Bi-directional cell-pericellular matrix interactions direct stem cell fate. Nature communications 9, 4049–12 (2018).

