## Supplementary data for "ILC1-derived TGFβ1 drives intestinal remodelling"

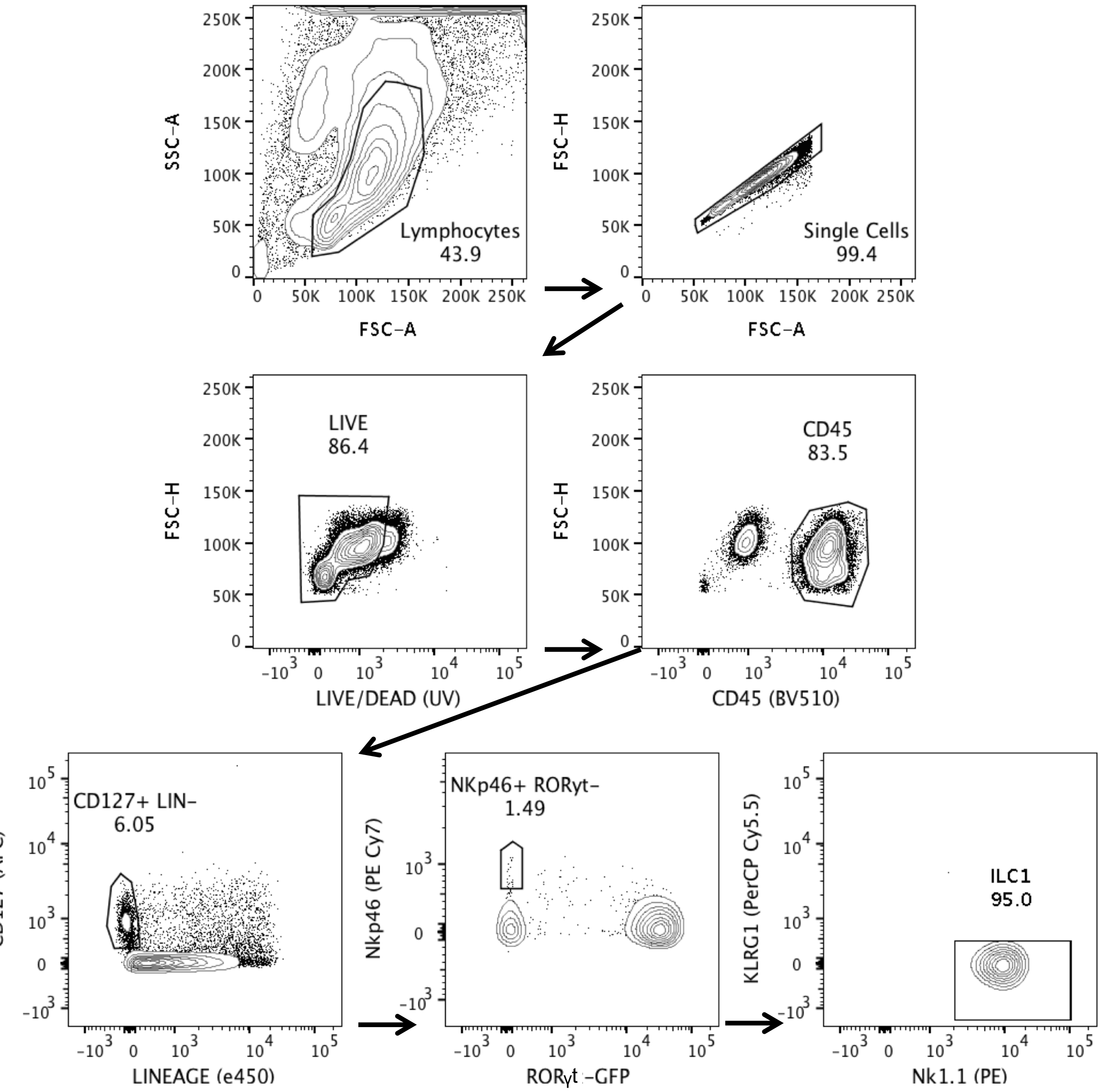


**Supplementary figure 1**. FACS gating strategy for the purification of ILC1 from the small intestinal lamina propria of female RORγt-GFP reporter mice

Gates were set based on relevant fluorescent minus one (FMO), used for Lineage (FITC), CD127 (APC), and NKp46 (PE Cy7). ILC1 were defined as single cells, live, $CD127^{+}$, Lineage (CD3, CD5, CD19, Ly6G)$^{-}$, RORγt$^{-}$, $NKp46^{+}$, $KLRG1^{-}$ and $NK1.1^{+}$.

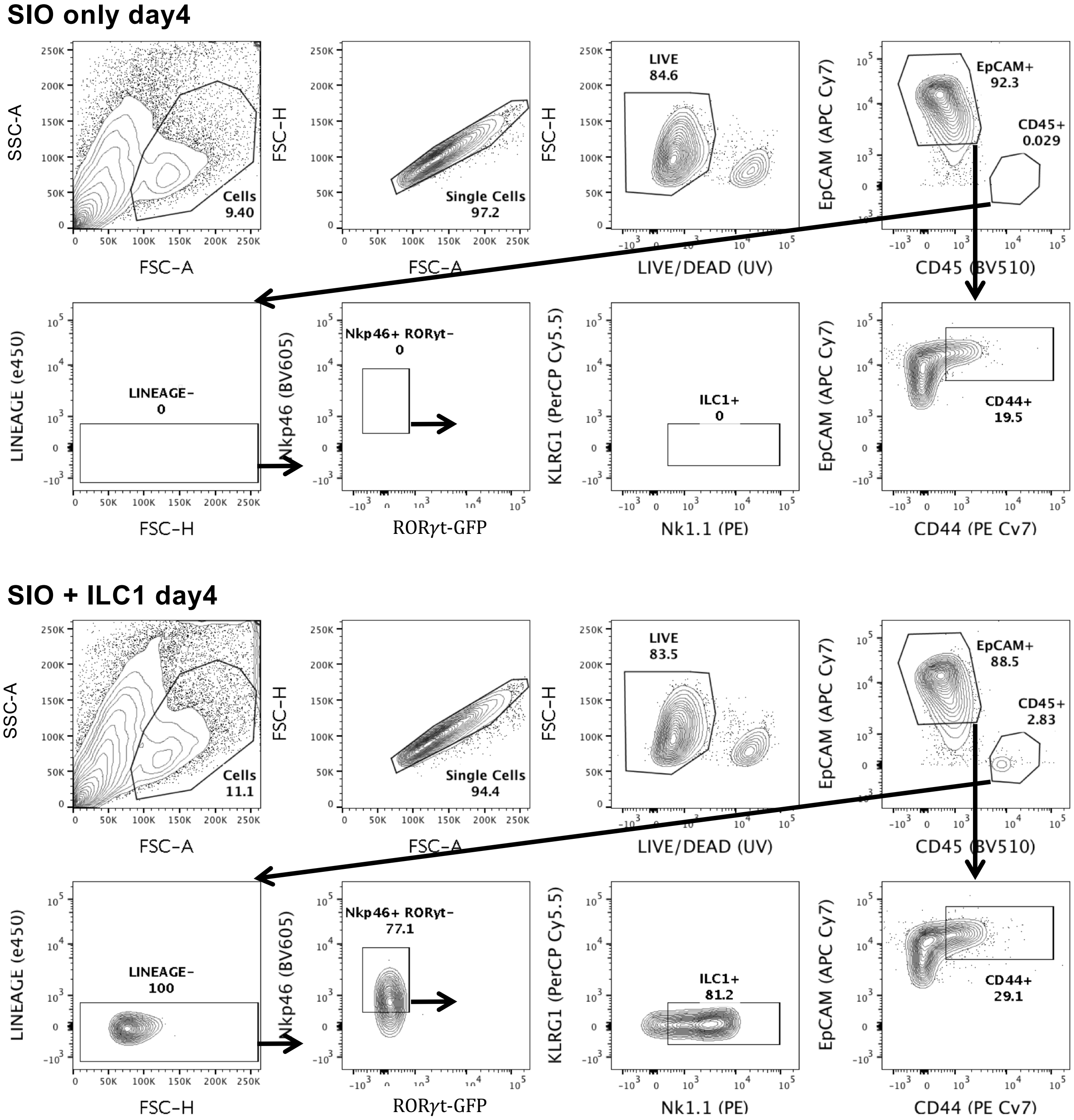


**Supplementary figure 2.** Gating strategy for FACS isolation of ILC1 and IEC from co-cultures

FACS gating strategy for dissociated ILC1-SIO co-cultures, sorting epithelial cells (single cells, live, $EpCAM^+$ and $CD45^-$) as well as ILC1 (single cell, live, $EpCAM^-$ $CD45^+$, Lineage (CD3, CD5, CD19, Ly6G)$^-$, $ROR\gamma t^-$, $NKp46^+$, $KLRG1^-$ and $NK1.1^+$). SIO only controls (top) were FACS purified to ensure all cells experienced the same conditions prior to analysis, and to allow for flow quantification of epithelial CD44 (IM7). Additional ILC markers were included in the panel to confirm no significant loss of ILC1, and that no contaminating non-ILC1 cells (e.g. Lineage positive cells or RORγt+ cells), or differences in ILC1 cell-count existed between samples. Gates were set on FMOs for CD44, Nkp46, and Lineage.

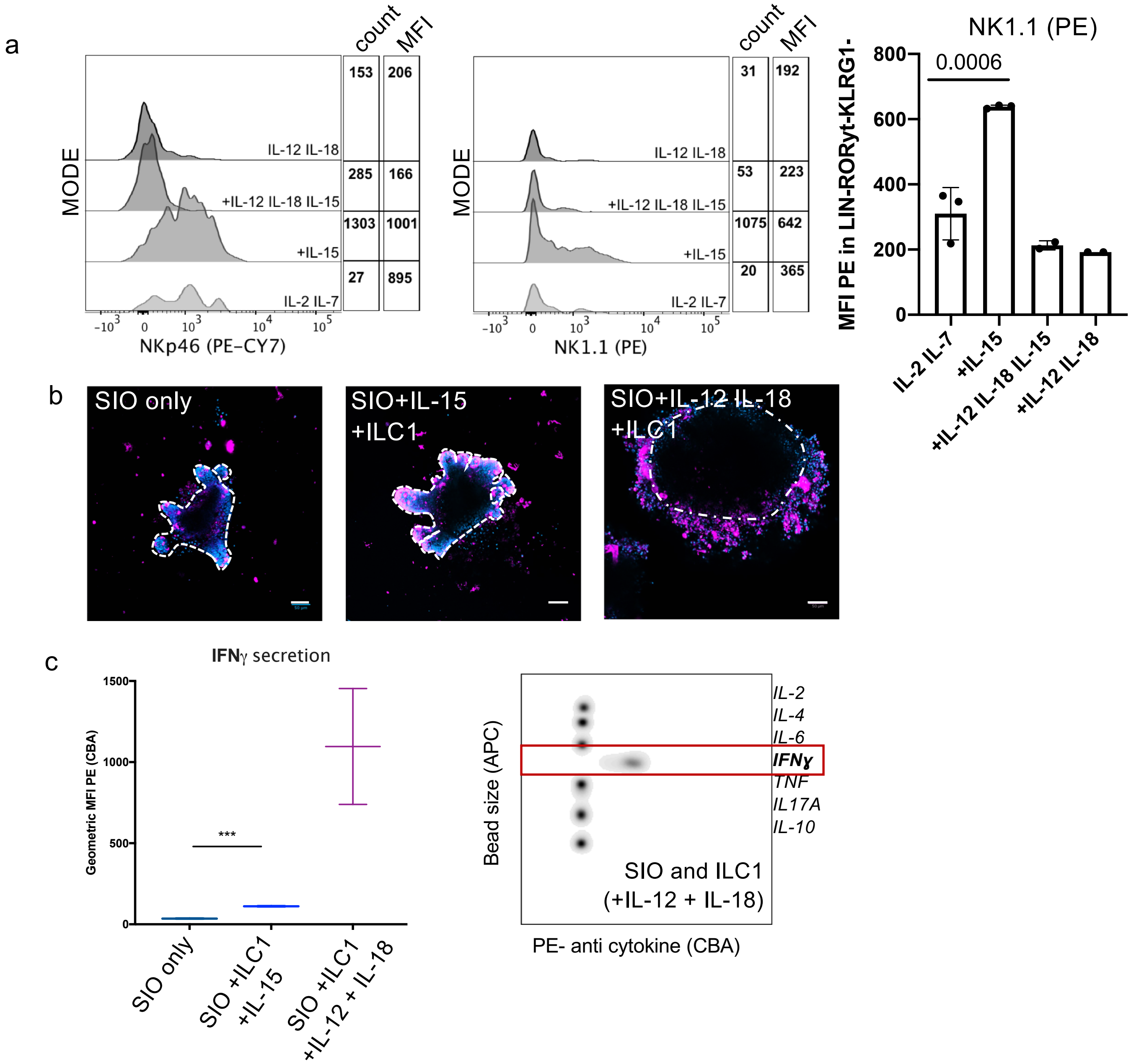


**Supplementary figure 3.** Optimization of ILC1-SIO co-culture conditions

a) Representative modal histograms of NKp46 and NK1.1 mean fluorescence intensity (MFI) from EpCAM-CD45+Lin-KLRG1-RORyt- ILC1 following 4 days of co-culture with SIOs supplemented with IL-2, IL-7 and a combination of ILC1 polarizing cytokines. IL-12 and IL-18 reportedly drive plasticity of ILC3 and ILC2 to ILC1. However, in co-culture with SIOs, addition of these cytokines reduced characteristic natural cytotoxicity receptor (NCR: NKp46) and NK1.1 expression. The measured mean fluorescent intensity of NK1.1 and ILC1 count was highest in conditions supplemented with 0.2ng/ml IL-15.

X-axis = Mode, representative of N=3, NK1.1 MFI quantified, OneWay ANOVA with Tukey's test shows significant difference between IL-2+IL-7 and IL-2+IL-7+IL-15 co-cultures, error bars S.E.M..

b) Representative confocal images of 4 day ILC1-SIO co-cultures supplemented with either IL-15 or IL-12 and IL-18 show dramatic loss of SIO morphology indicative of potential loss in SIO viability in conditions with IL-12 and IL-18 with ILC1 (right). Scale bar 50µm.

c) Addition of IL-15 induces significant secretion of IFNγ. IL-12 and IL-18 increase IFNy expression, but not of other Th1/Th2/Th17 cytokines, as measured by flow Cytometric Bead Array (CBA), in which supernatant of co-culture was stained with APC-bead conjugated antibodies of different sizes, and then counterstained with PE-conjugated antibodies against cytokines, a method similar to an ELISA. Representative of N=2, error bars S.D.

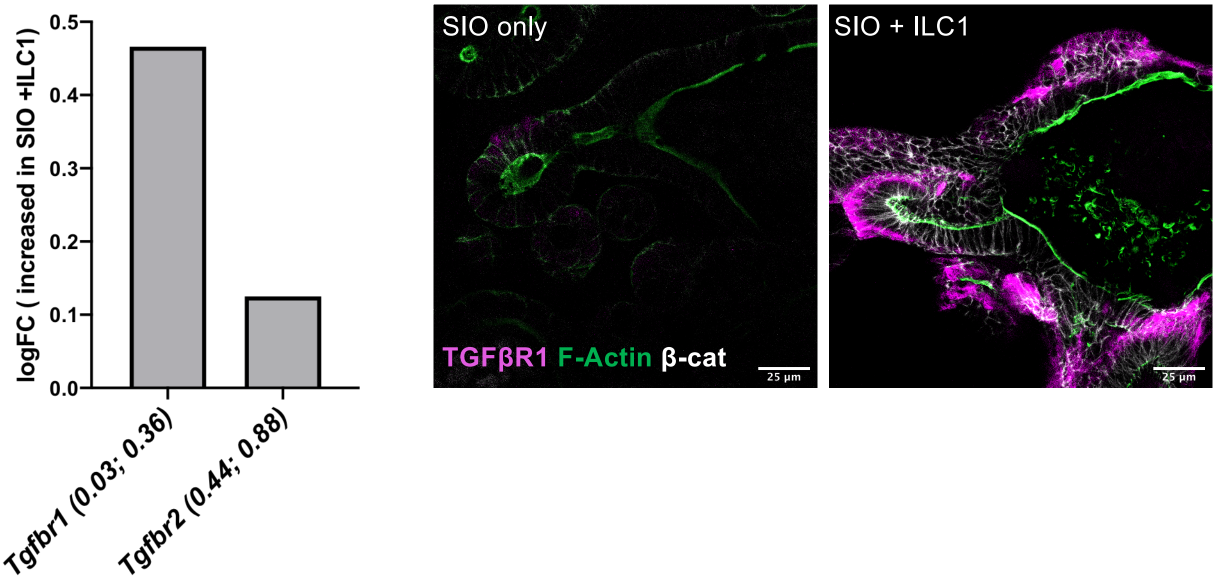


**Supplementary figure 4.** Differential upregulation of TGFβR1 expression in SIO after co-culture with ILC1

a. LogFC of TGFβ Receptors in SIO after co-culture with ILC1 from initial RNAsequencing dataset (N=3, brackets indicate p value and padj). Representative confocal image of SIO only and SIO+ILC1 showing TGF**R1** expression in magenta, F-actin in green, and β-catenin in white (Rep of N=2, scale bar 25$\mu$m.)

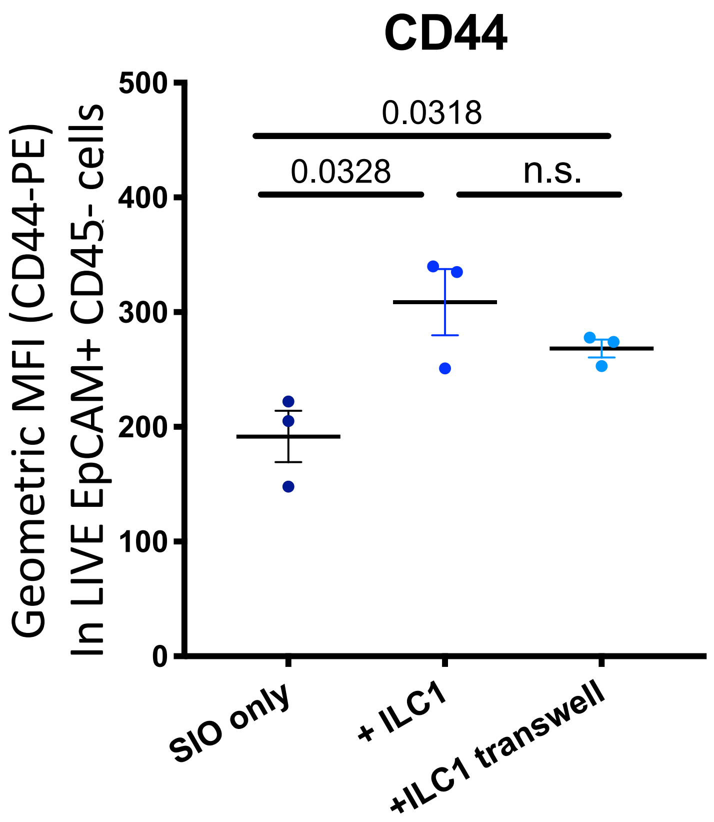


**Supplementary figure 5.** ILC1 impact on IEC is not contact dependent
ILC1 separated from SIO by a transwell insert still drive significant upregulation of CD44 (IM7) MFI, as measured by flow cytometry in LIVE EpCAM+ CD45- cells. OneWay Anova with Tukey's test, N=3, error bars S.E.M.

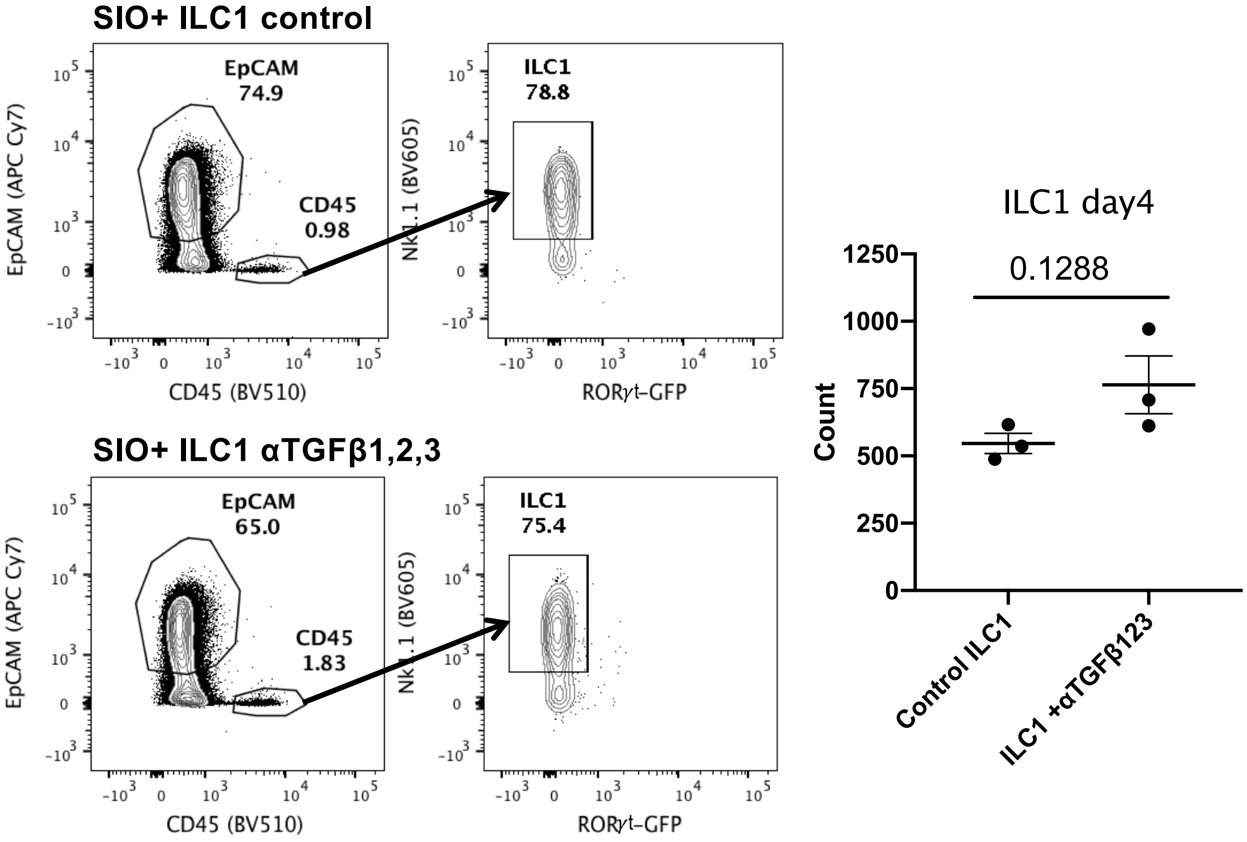


**Supplementary figure 6.** TGFβ1,2,3 neutralization in ILC1-SIO cultures does not impact ILC1 phenotype

Representative plots from 4 day ILC1-SIO co-cultures with or without TGFβ1,2,3 neutralization (500 ng/ml) show that percentage of ILC1 (as defined in Supplementary Fig. 2) and count of LIVE, EpCAM-, CD45+, Nkp46+, RORγt-, NK1.1+ ILC1 (ILC1 from N=3 individual mice) do not change significantly (Unpaired two-tailed student t-test), suggesting that any decrease in CD44 expression is not due to a loss of ILC1 phenotype or survival.

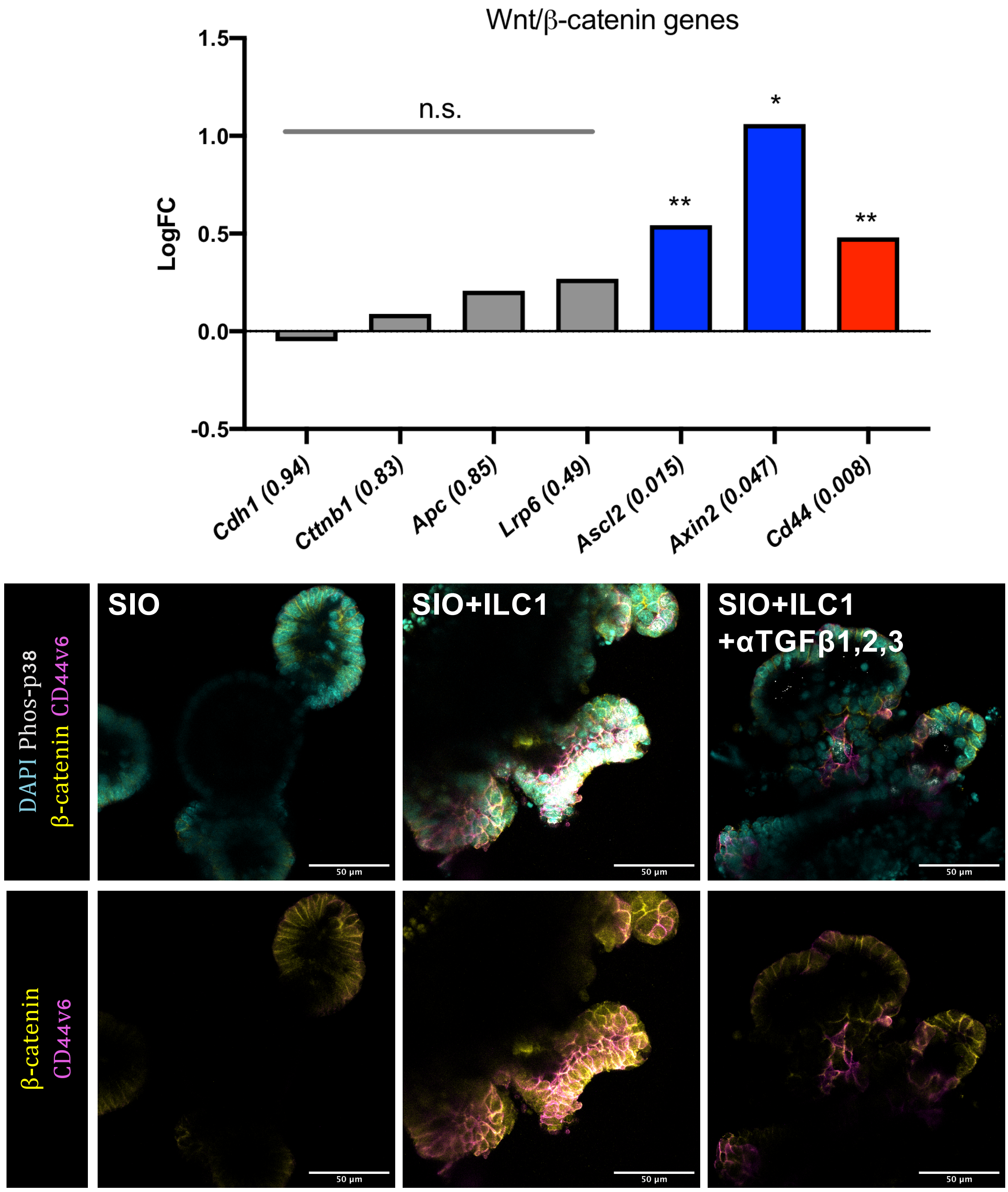


**Supplementary figure 7.** ILC1 increase epithelial β-catenin expression

a. LogFC expression of key genes involved in Wnt/β-catenin signaling in SIO + ILC1 cultures relative SIO only controls on day4, extracted from the RNAsequencing dataset. Brackets indicate padj values, with n.s. non-significantly differentially regulated genes represented by grey boxes (N=3).

b. Additional representative images relating to Fig. 2g-i, highliting ILC1 induced, TFGβ1-dependent β-catenin accumulation across the basolateral membrane and cytoplasm, which appear to overlap with CD44v6 expression (bottom). Scale bars 50$\mu$m.

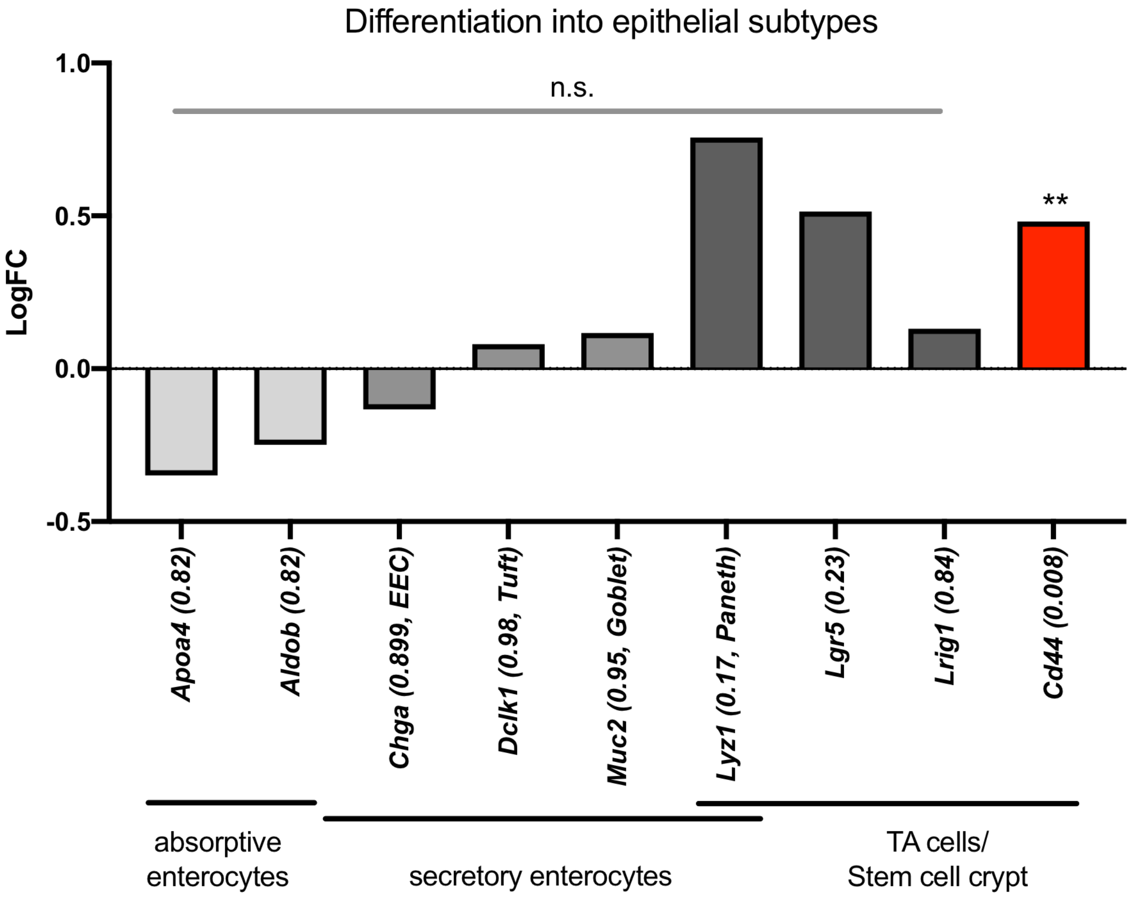


**Supplementary figure 8.** ILC1 do not significantly alter subset specific gene expression

a. LogFc of differentially expressed genes in SIO+ILC1 relative to SIO only on day4, extracted from RNA-sequencing dataset (N=3) show no significant difference in key marker expression relative to SIO only control. padj values indicated in brackets after gene names, along with the epithelial subtype they characterise.

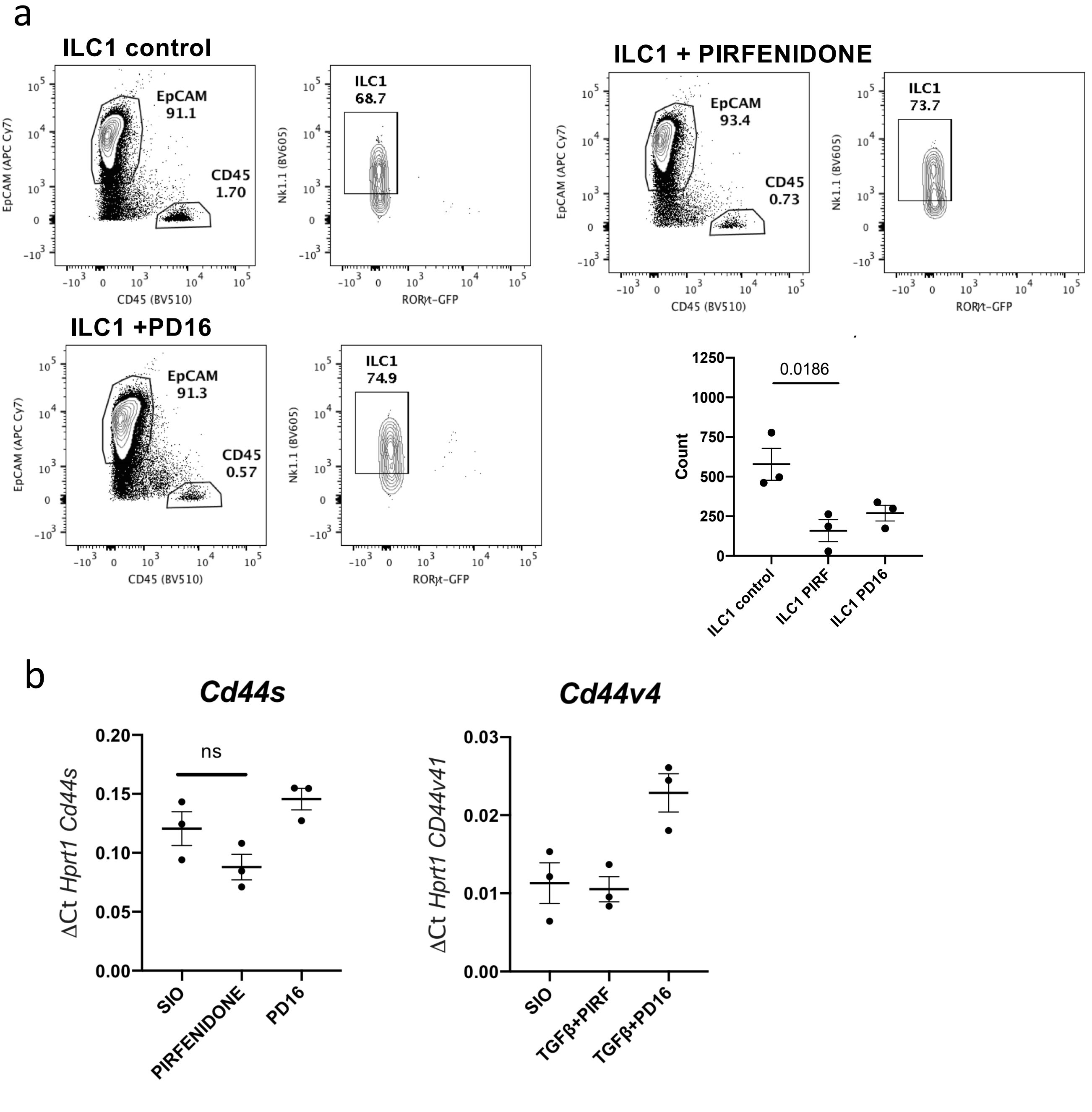


**Supplementary figure 9.** p38 inhibition decreases number of ILC1 in co-culture

a. Representative FACS plots from 4 day ILC1-SIO co-cultures with regular IL-15 media or supplemented with Pirfenidone (50$\mu$M) or PD16 (5$\mu$M). While ILC1 phenotype itself was not significantly impacted, the number of ILC1 (count live, EpCAM- CD45+) after co-culture decreased significantly to below 250 cells/50000 recorded events (~2000 cells per condition). Count: OneWay ANOVA with Tukey's test of ILC1 co-cultures from N=3 mice. Error bars S.E.M.

b. RTqPCR of *Cd44s* and *Cd44v4* in SIO cultured alone, with Pirfenidone, or PD16 for 4days. Two-tailed student t-test between SIO and Pirfenidone in *Cd44s* condition show no significant difference in *Cd44s* or *Cd44v4* expression induced by Pirfenidone; Error bars S.E.M.

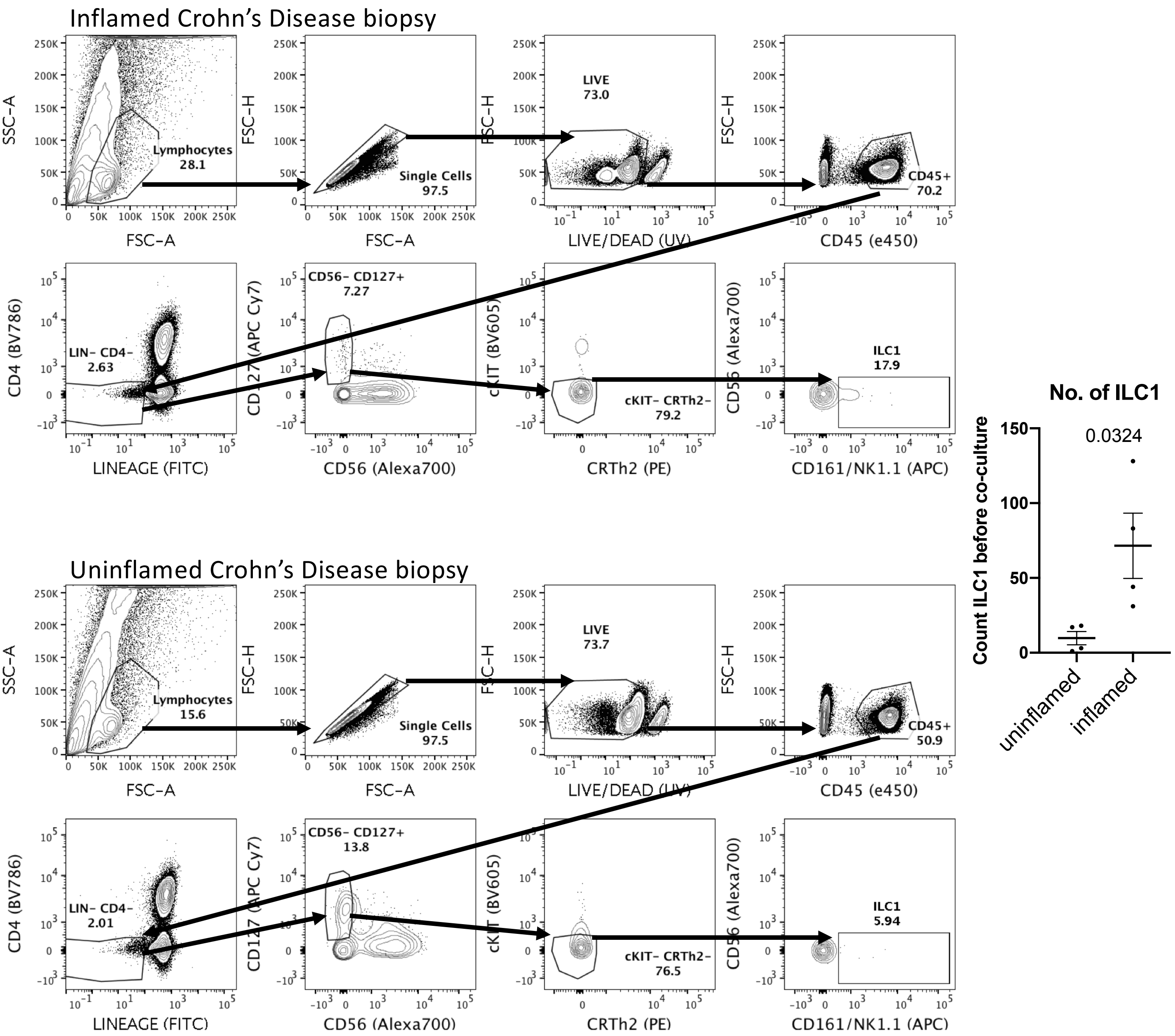


**Supplementary figure 10.** Gating strategy for hILC1 from patient lamina propria biopsies

Colonic lamina propria ILC1 were purified from 15-18 biopsies of IBD (predominantly Crohn's Disease) patients with or without active inflammation. hILC1 were defined as LIVE, CD45+, CD4-, Lineage- (CD3, CD4, CD14, CD19, CD20, TCRαβ, TCR$\gamma\delta$), CD127+, CD56- (NK cells), cKIT- (ILC3 and precursors), CRTh2- (ILC2), CD161/Nk1.1+. FACS plots representative of variation in hILC1 yield between patients, with a significant difference in ILC1 count from inflamed versus uninflamed (count of representative 100,000 events recorded during FACS, N=6, unpaired two-tailed student t-test, error bars S.E.M.)

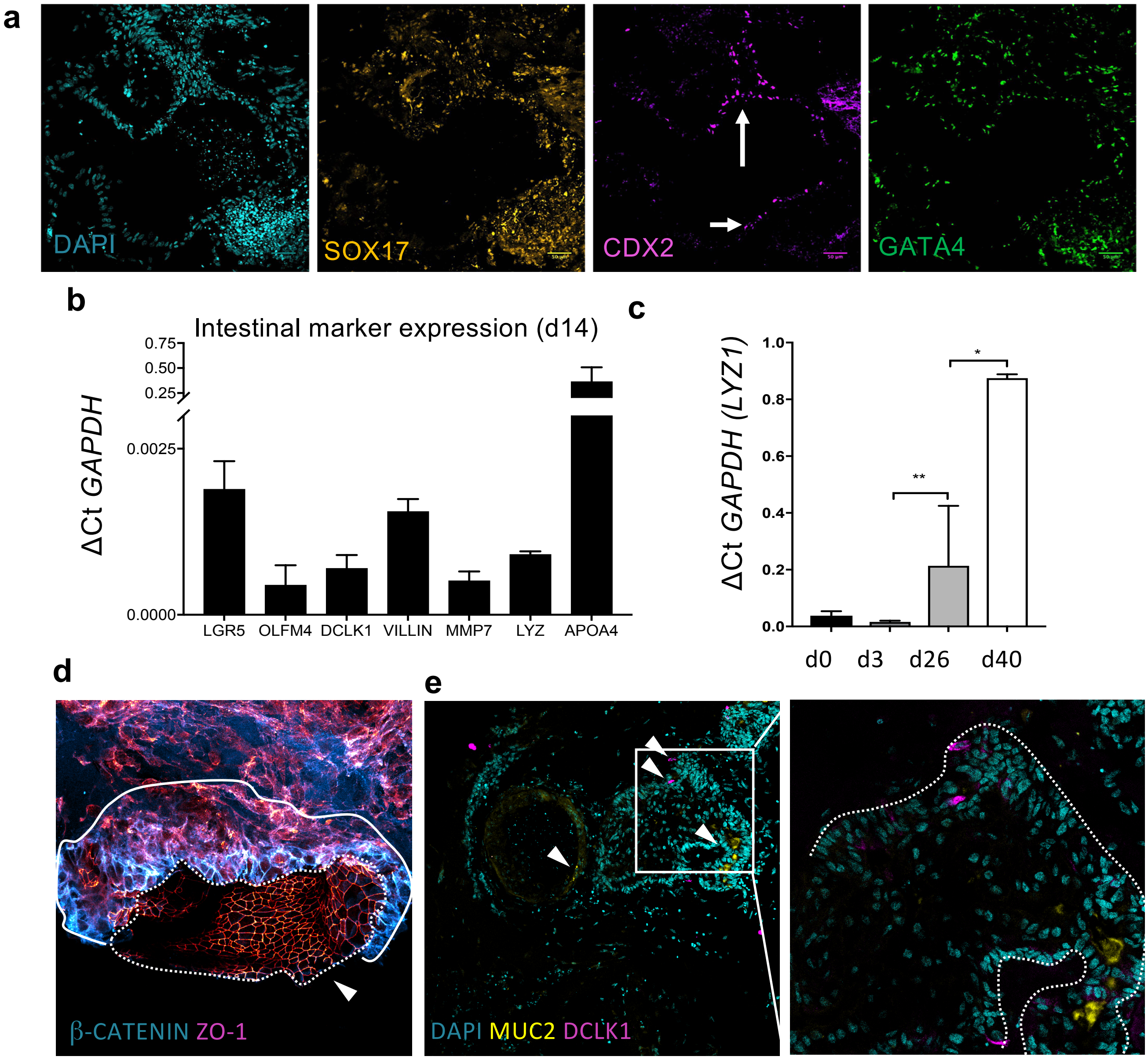


**Supplementary figure 11.** Characterisation of HIO development in Matrigel )

a. Representative confocal images of 10 day old HIOs demonstrating characteristic definitive endoderm markers, with CDX2 expression restricted to the future epithelial buds that are picked and cultured in 3D Matrigel. Scale bar = 50$\mu$m. b. HIO structures on day 14 express characteristic epithelial markers after 14 days of differentiation. Error bars represent S.D., N=2 differentiations. c. HIO increase expression of anti-microbial Paneth cell marker Lyzozyme1 over the time course of differentiation (N=2, error bars S.D., unpaired student t-tests between d3-d26, and d26-d40). d. Correct apico-basal expression of apical tight junction marker ZO-1 facing into the pseudolumen (indicated by white arrow, outlined in white dotted line), with β expression indicating organization in an epithelial structure in a d75 HIO. e. Rep. confocal image and enhanced zoom of white box show that d75 HIO express markers of mature intestinal epithelium like Stem and Tuft cell marker DCLK1 (top white arrows) and deposition of mucous layer component MUC2 into the pseudolumen.

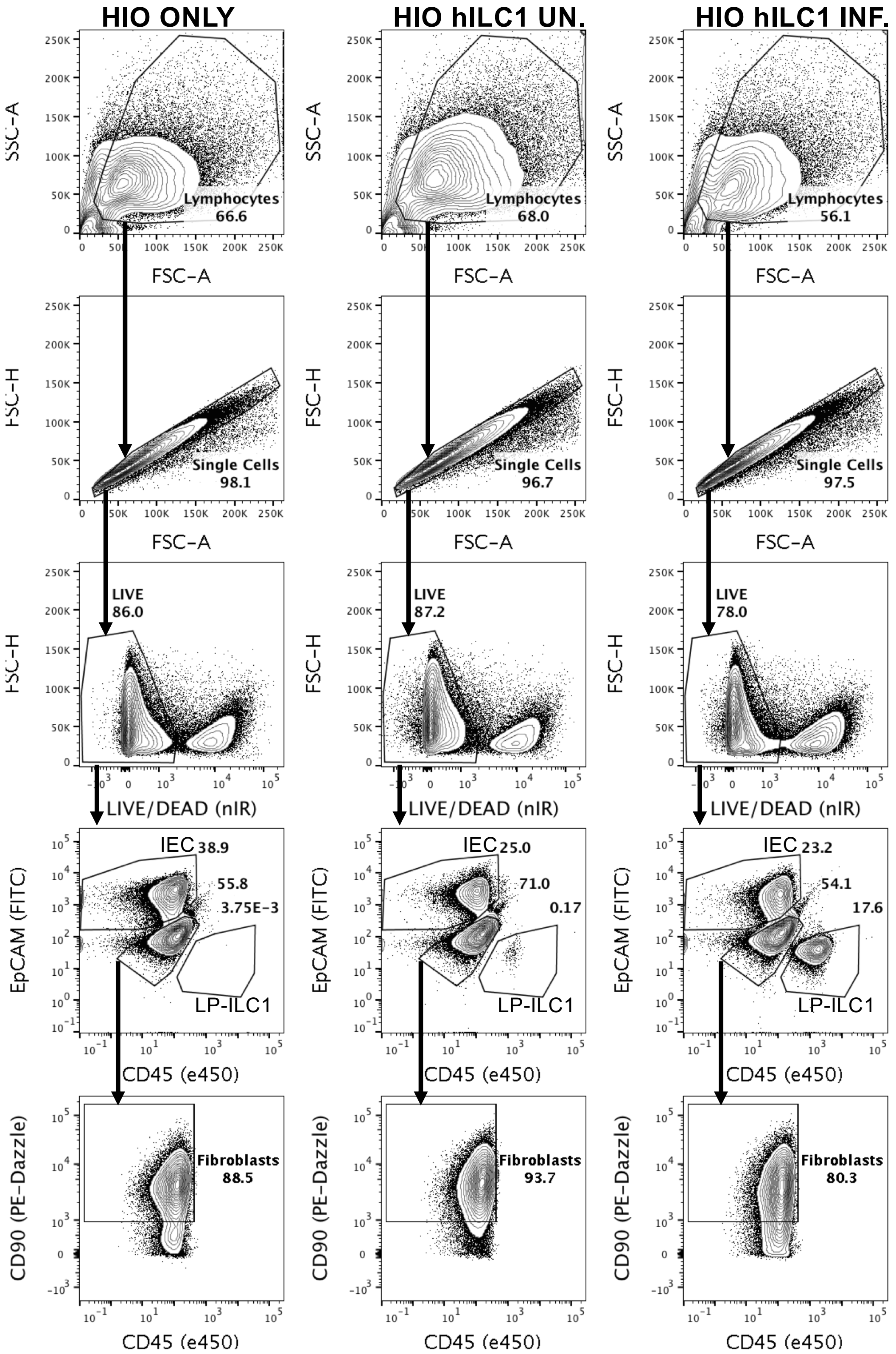


**Supplementary figure 12.** HIO and hILC1 cell subtype FACS gating strategy after 7 day co-culture Epithelial cells from co-cultures were defined as single, live and EpCAM$^{+}$. Fibroblasts were defined as single cells, live, EpCAM$^{-}$, CD45$^{-}$ and CD90$^{+}$. hILC1 were defined as single cells, live, EpCAM$^{-}$, CD45$^{+}$

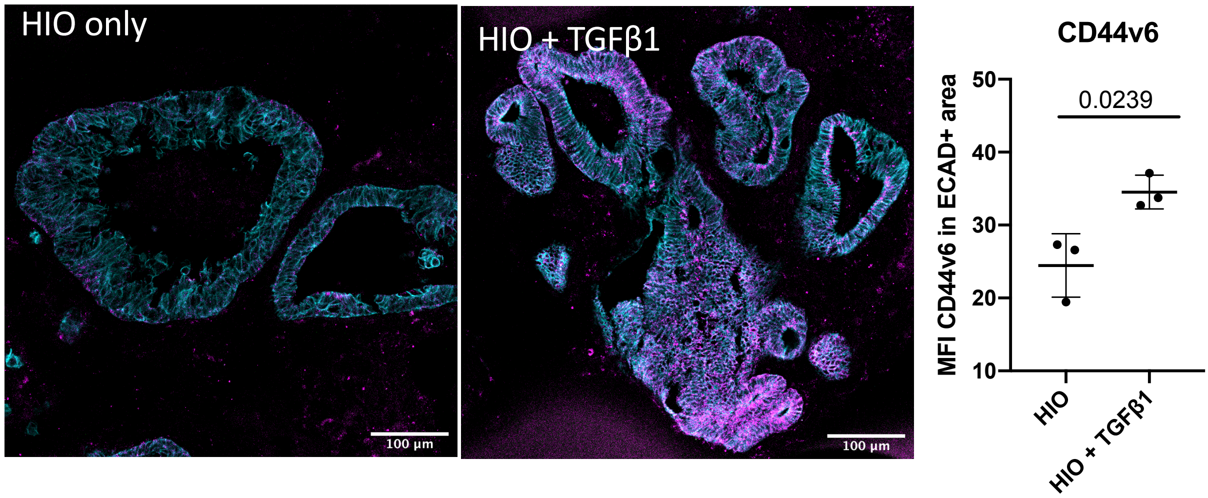


**Supplementary figure 13** TGFβ1 induces expression of CD44v6 in HIO

Representative confocal images CD44v6 (magenta) expression in HIO and HIO after 7 day co-culture with recombinant TGFβ1, with MFI of CD44v6 quantified in E-cadherin (cyan) regions (N=3). Scale bars 100$\mu$m. Unpaired student t-test, error bars S.E.M.

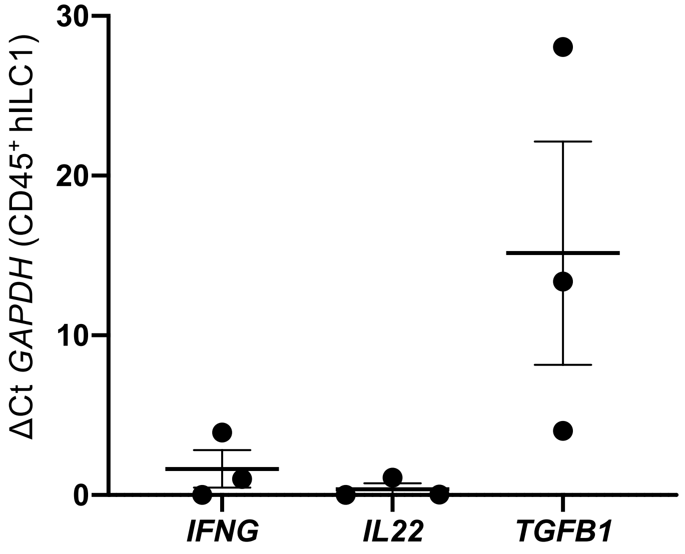


**Supplementary figure 14.** hILC1 from inflamed IBD patients express TGFB1
Expression of *IFNG*, *IL22*, and *TGFB1* from activated uninflamed (N=1) and inflamed (N=2) patient biopsy hILC1.

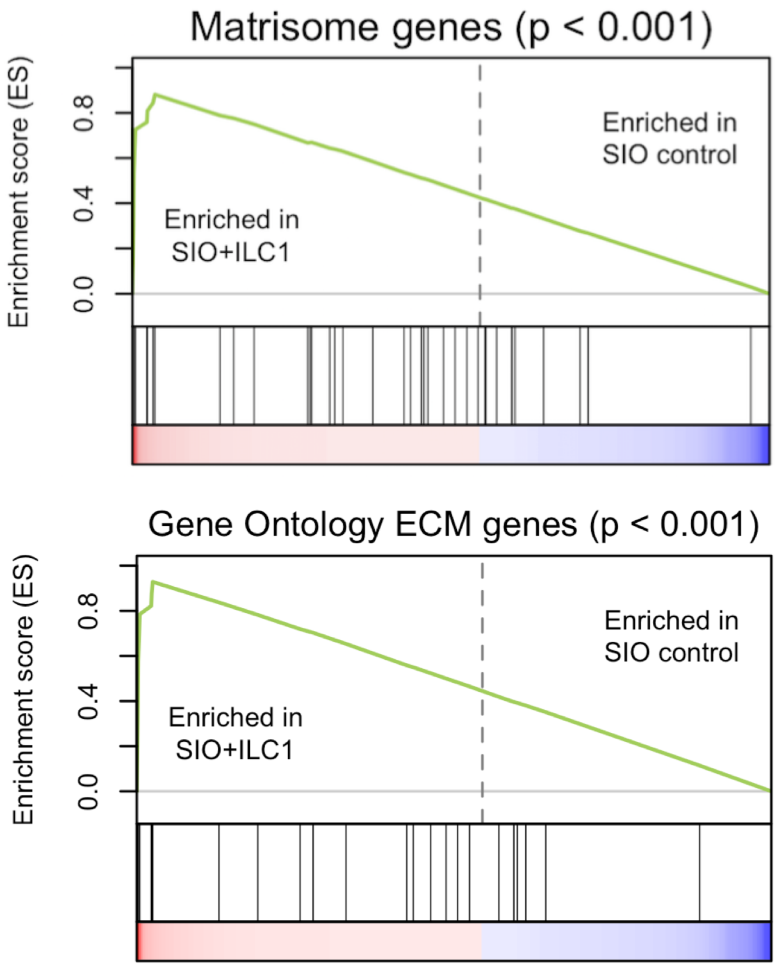


**Supplementary figure 15.** GSEA analysis of murine RNA-sequencing shows enrichment of matrisome and ECM-associated genes in SIO co-cultured with ILC1 (N=3).

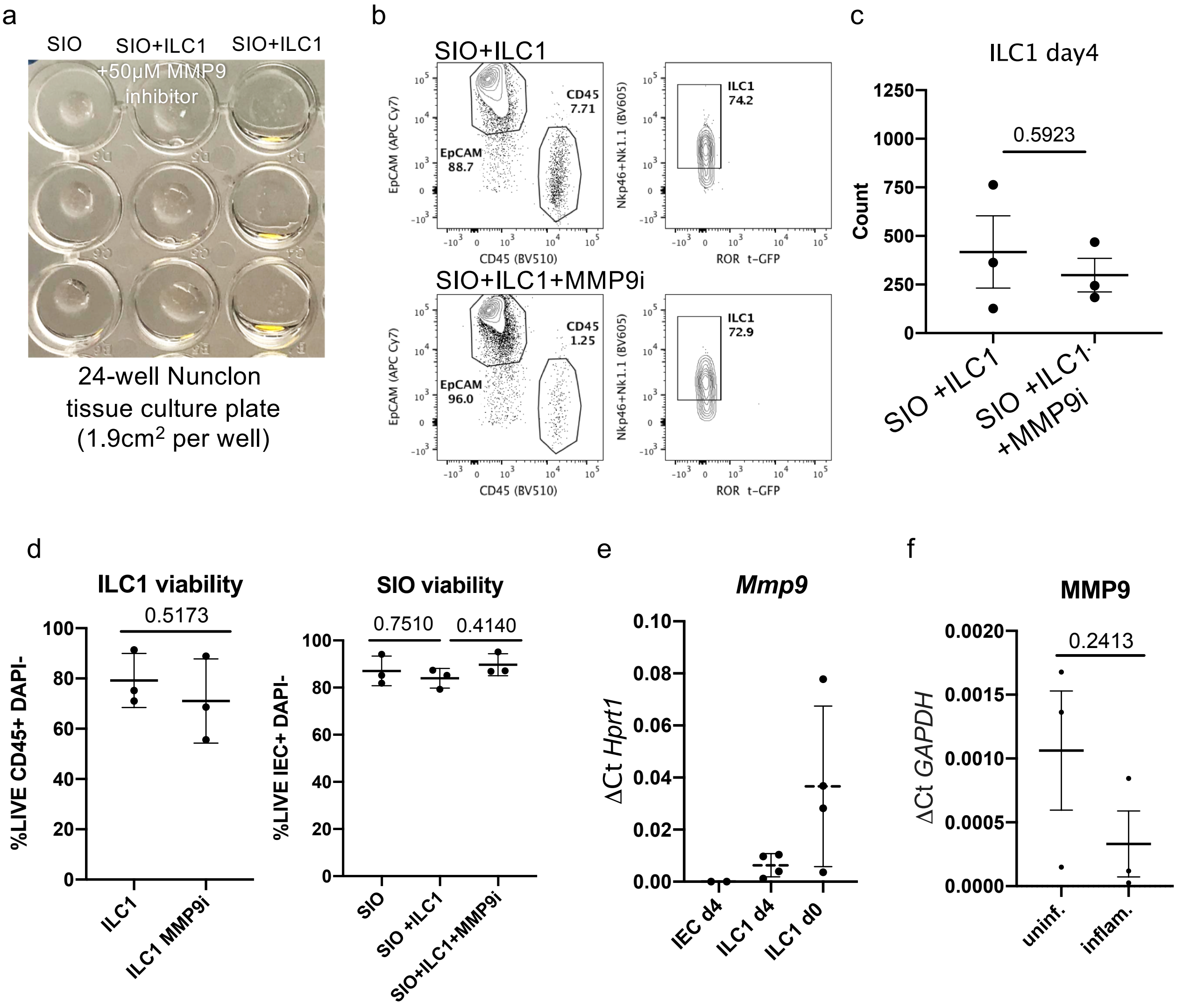


**Supplementary figure 16.** ILC1 express MMP9 and drive Matrigel degradation

a) Image of Matrigel co-culture integrity after 24h with SIO only, SIO+ILC1+MMP9 specific inhibition (MMP9i), or SIO+ILC1 show complete degradation of Matrigel bubble in ILC1 co-cultures, reversible by MMP9i. SIO in SIO+ILC1 condition are pooled in PBS at the bottom of the well, or are loosely attached to the tissue culture plastic (ILC1 from N=3 mice).

b) Representative plots from day 4 ILC1-SIO co-cultures with MMP9 inhibition (50$\mu$M) show (c) no significant loss of ILC1 cell count due to MMP9i.

d) Flow cytometry analysis of viability (DAPI) in CD45+EpCAM- ILC1 (left) and CD45-EpCAM+ IEC on day4 of co-culture with or without MMP9i.

e) *Mmp9* expression in murine ILC1 before (N=4, d0) and after (N=3, d4) co-culture with SIO, and in CD45-EpCAM+ IEC after co-culture.

f) Relative expression of *MMP9* in human hILC1 from uninflamed or inflamed patients after 7 day co-culture with HIO. Unpaired two-tailed t-test, error bars S.E.M.

**a**

| | System | PEG-4NPC | PEG-4VS | Peptides | Ions | Water |
|---|---|---|---|---|---|---|
| $A_4$+$B_4$ | Ac-KDWERC-NH2 | 40 | 40 | 0 | 160 $Na^+$ | 533336 |
| | H-SREWERC-NH2 | 40 | 40 | 0 | 0 | 533336 |
| $A_2$+$B_4$ | Ac-CREWERC-NH2 | 0 | 80 | 160 | 0 | 533336 |

**b**

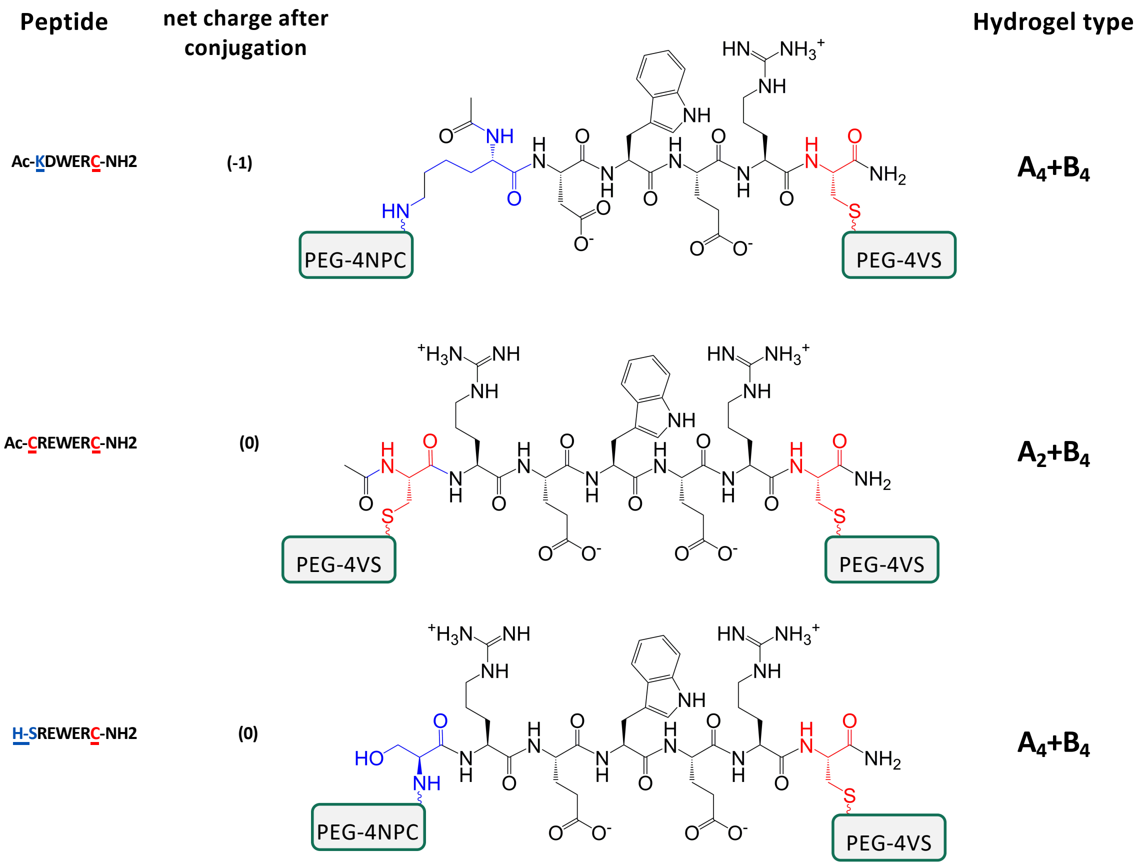


**Supplementary figure 17**. Hydrogel systems simulated by molecular dynamics.
a) Table outlining the systems characterized using molecular dynamics simulations with details on the number of pre-conjugated PEG-peptide molecules (PEG-4NPC), PEG molecules (PEG-4VS), free peptides, and ions in each simulation. The number of water beads was identical in each system and is 4 times the number of beads. Systems are designated by their cross-linking peptide. Simulations were run twice for each condition.
b) Chemical structure of peptides used in simulations. Peptide Ac-KDWERC-NH2was the non-adhesive/non-degradable peptide in our experimental hydrogel design ($A_4$+$B_4$). Simulations were also run with ‘mutated’ peptides H-SREWERC-NH2 and Ac-CREWERC-NH2.

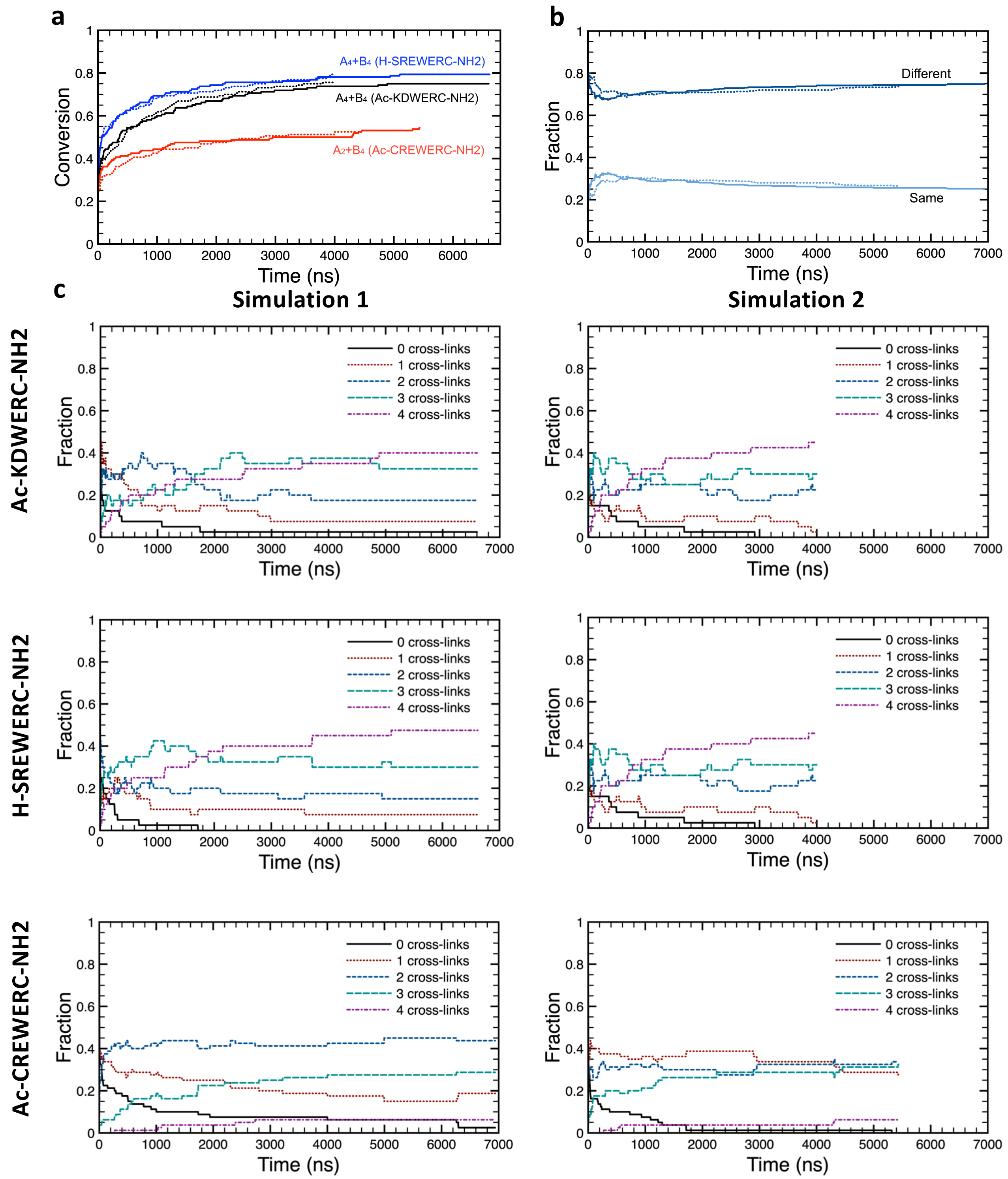


**Supplementary figure 18**. Additional findings from molecular dynamics simulations.
a) Fraction of total possible network forming bonds that form in simulated systems. Lines show two independent simulations per condition. The H-SREWERC-NH2 peptide forms 0.79 of possible network forming bonds.
b) Fraction of peptides in the $A_2$+$B_4$ design (Ac-CREWERC-NH2) that form two new bonds between arms of the *same* PEG molecule (primary loops) or between two *different* PEG molecules (cross-linking). These observations are in line with reported experimental observations of 1° loop formation in $A_2$+$B_4$ systems (Gu, Y. *et al., PNAS (*2017)). Bonds between arms of the same PEG molecule are precluded in simulations of the $A_4$+$B_4$ design.
c) Fraction of PEG molecules (PEG-4VS) that have formed 0, 1, 2, 3, or 4 network forming bonds (cross-links) as a function of time for the 3 different systems. As ~25% of bonds formed in the $A_2$+$B_4$ design (Ac-CREWERC-NH2) are not network forming, a larger fraction of PEG molecules form only 1 or 2 network forming cross-links, as opposed to the $A_4$+$B_4$ designs where the majority of PEG molecules form 3 or 4 network forming cross-links.

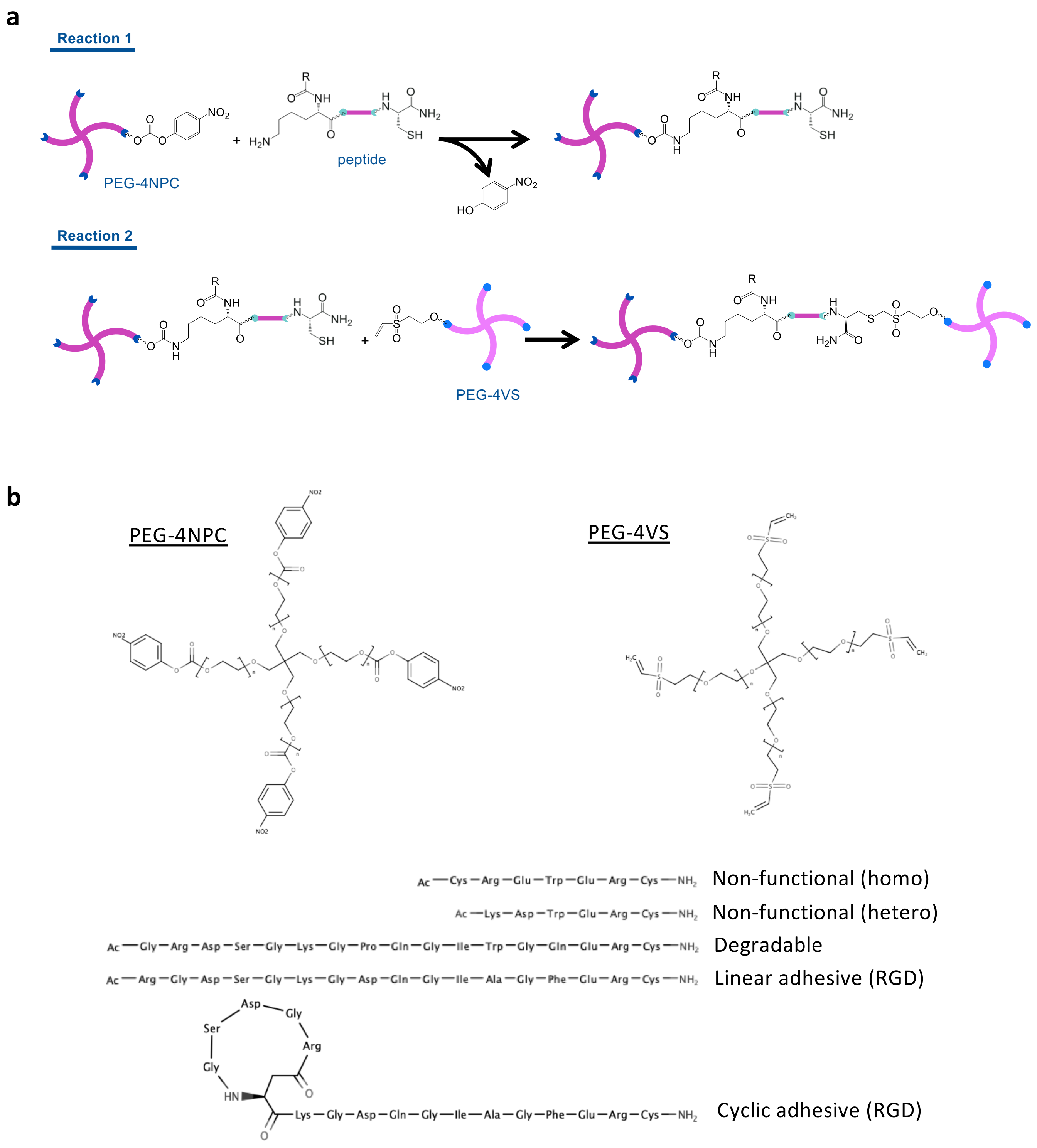


**Supplementary figure 19**. Chemical strategy to form PEG-based hydrogels.
a) Reaction schemes used to form hydrogels.
b) Chemical structures of PEGs and peptides used to form hydrogels. During peptide synthesis, the N-terminal amine was acetylated (or reacted with an aspartate side chain for cyclic adhesive RGD), rendering it non-reactive. Peptides were designed such that the degradable sequence is placed in between the peptide cross-linking points, i.e. between the lysine and cysteine residues, while the RGDS sequences are placed at the N-terminal and before the reactive lysines. In many $A_2$+$B_4$/$B_8$ designs, cell adhesive peptides are presented through a pendant chemical group that does not contribute to cross-linking, meaning that modulation of ligand concentration may result in concurrent unintended modulation of cross-linking and consequently stiffness. By rendering all peptides capable of cross-linking, our design ensures that ligand density can be modulated independently of stiffness.

**a**

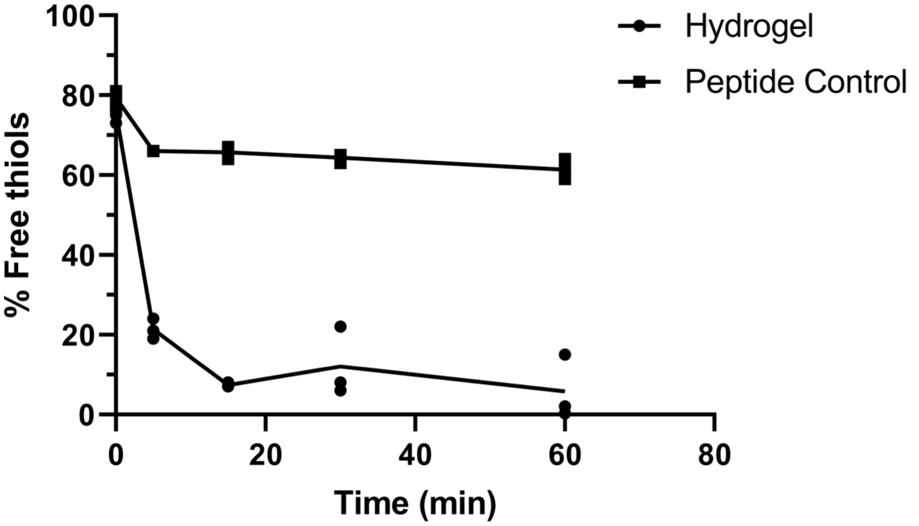


**b**

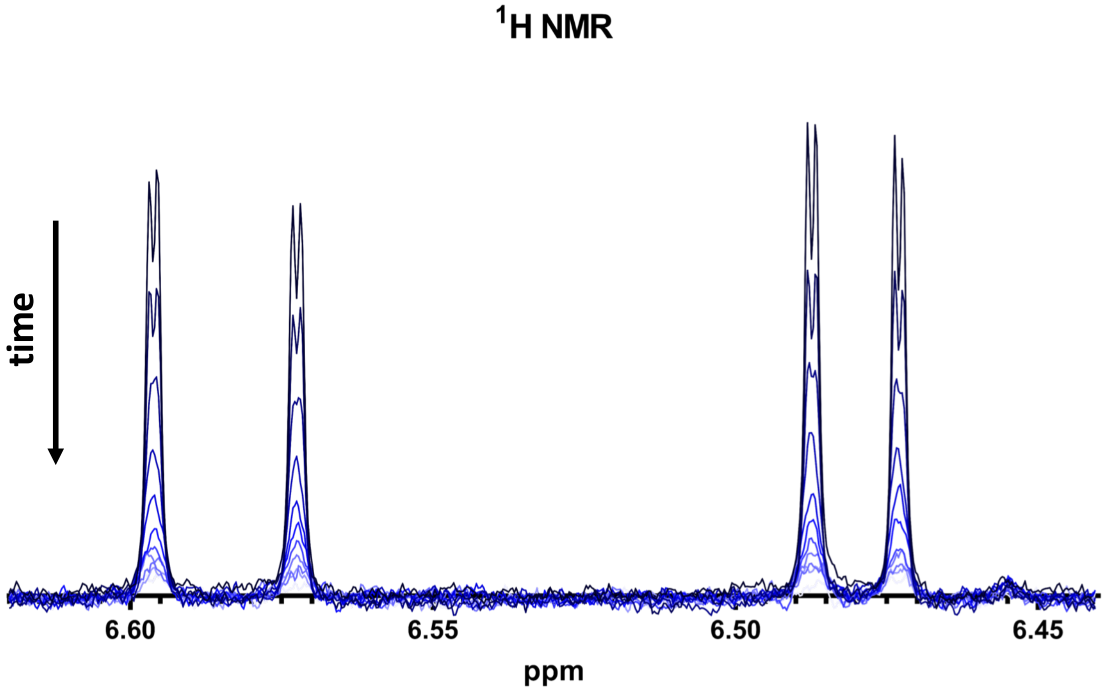


**Supplementary figure 20**. Analysis of the efficiency of the click reaction used to form hydrogels.
a) Plot showing the percent of total free thiols consumed during the Michael addition reaction to form hydrogels as determined by Ellman's assay. Nearly 80% of free thiols were consumed within 5 min and 94% within 1 hr. Lines connect mean values (N=3).
b) Proton NMR spectra showing the kinetics of the reaction over 60 min by monitoring the disappearance of olefinic vinyl bonds at 6.5841 (M=2, J1= 16.6000 Hz, J2= 0.8959 Hz) and 6.4800 (M=2, J1= 10.0440 Hz, J2= 0.9131 Hz) ppm as a result of the Michael addition. Within 6 min, 70% of the vinyl groups had disappeared, and 99% were gone after 60 min. Lines are coloured lighter as time proceeds.

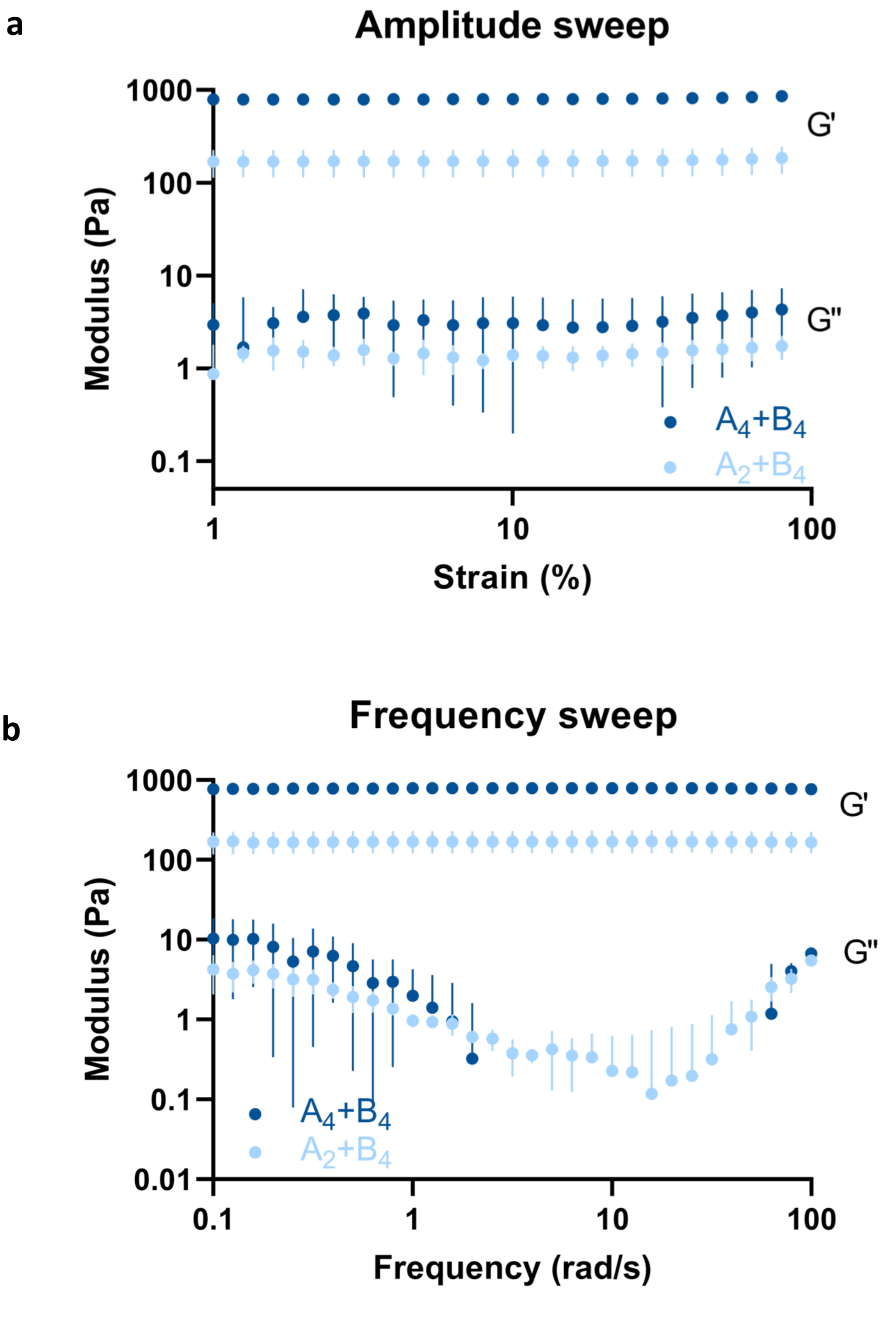


**Supplementary figure 21**. Rheological characterization of $A_4$+$B_4$ and $A_2$+$B_4$ hydrogels
a) Mean (± standard deviations) values of G' and G'' obtained using amplitude sweeps on $A_4$+$B_4$ and $A_2$+$B_4$ hydrogels (N=3).
b) Mean (± standard deviations) values of G' and G'' obtained using frequency sweeps on $A_4$+$B_4$ and $A_2$+$B_4$ hydrogels (N=3).
For some values of strain and frequency, G'' for $A_4$+$B_4$ hydrogels could not be obtained. Taken together with the time sweep measurements (Fig. 4e), these data suggest that $A_4$+$B_4$ hydrogels have a higher elastic modulus, behave more elastically, and are more reproducible (less inter-sample variability) than $A_2$+$B_4$ designs.

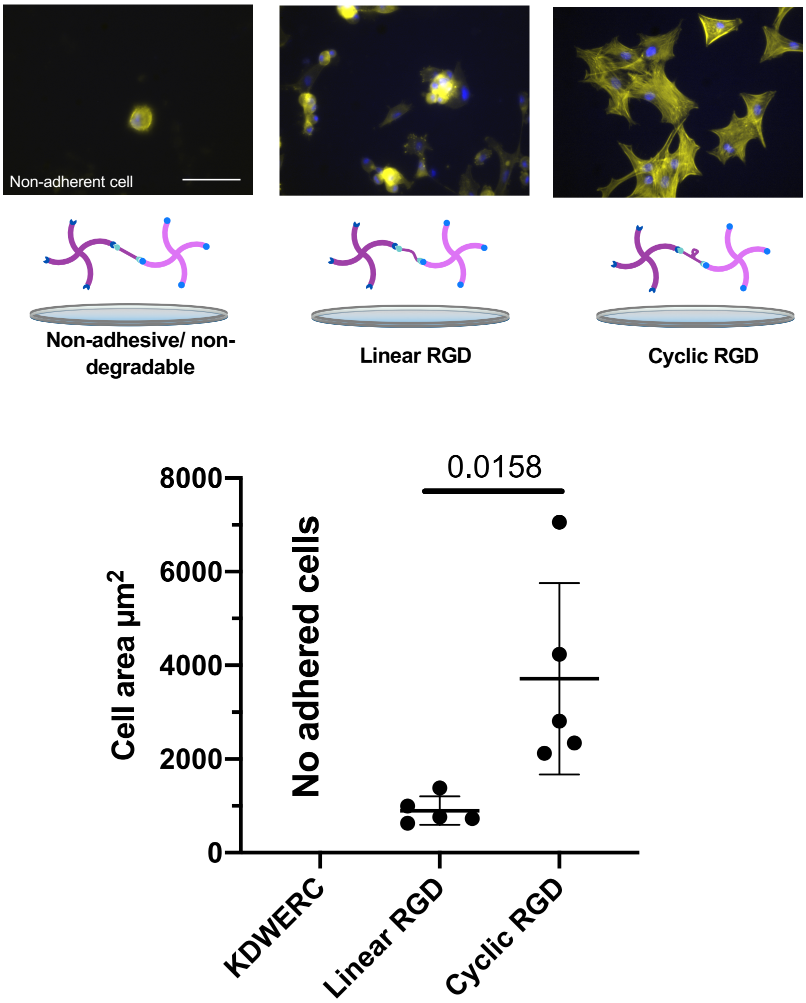


**Supplementary figure 22**. hMSC adherence on 2D hydrogel surfaces.
Representative fluorescence micrographs of hMSC cultured on 2D hydrogels stained with phalloidin-TRITC and DAPI. hMSC do not adhere to hydrogels formed with non-adhesive/non-degradable peptides. On hydrogels formed with a linear RGD sequence, hMSC display a mixture of round and spread morphologies, but when all peptides contain a cyclic RGD sequence, hMSC adopt highly spread morphologies. Unpaired student t-test between Linear RGD and cyclic RGD, error bars S.E.M. Scale bar = 50$\mu$m.

a

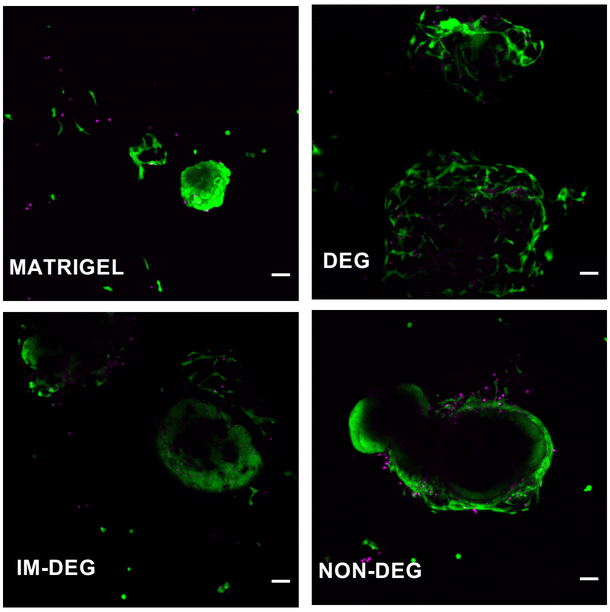


b

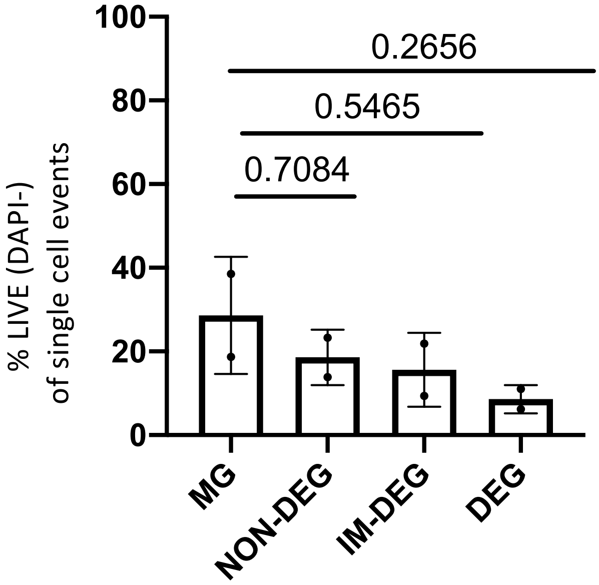


**Supplementary figure 23**. Cytotoxicity assay of HIO in hydrogels

a) Representative live/dead images (FDA/Propidium Iodide) from HIO encapsulated in Matrigel, in 75%degradable(DEG), 45%degradable (IM-DEG), and 0%degradation (NON-DEG) hydrogesl. Scale bar 50$\mu$m (N=2). b) Flow cytometry of HIO in Matrigel (MG), or encapsulated in NON-DEG, IM-DEG, or DEG hydrogels show no significant differences in viability between Matrigel and any hydrogel condition. Viability appears lower (~20%) relative to live/dead stainings (~80%) due to the single cell dissociation and prolonged trypsin-treatments necessary to retrieve HIO from crosslinked hydrogel, however values are representative of the lack of statistically significant differences in viability between conditions, which were dissociated via the same technique (N=2 technical replicates, unpaired two-tailed student t-test between MG and each condition).

FEEL FREE TO DELETE THIS PART, EDIT AND SWITCH BACK TO BLACK.

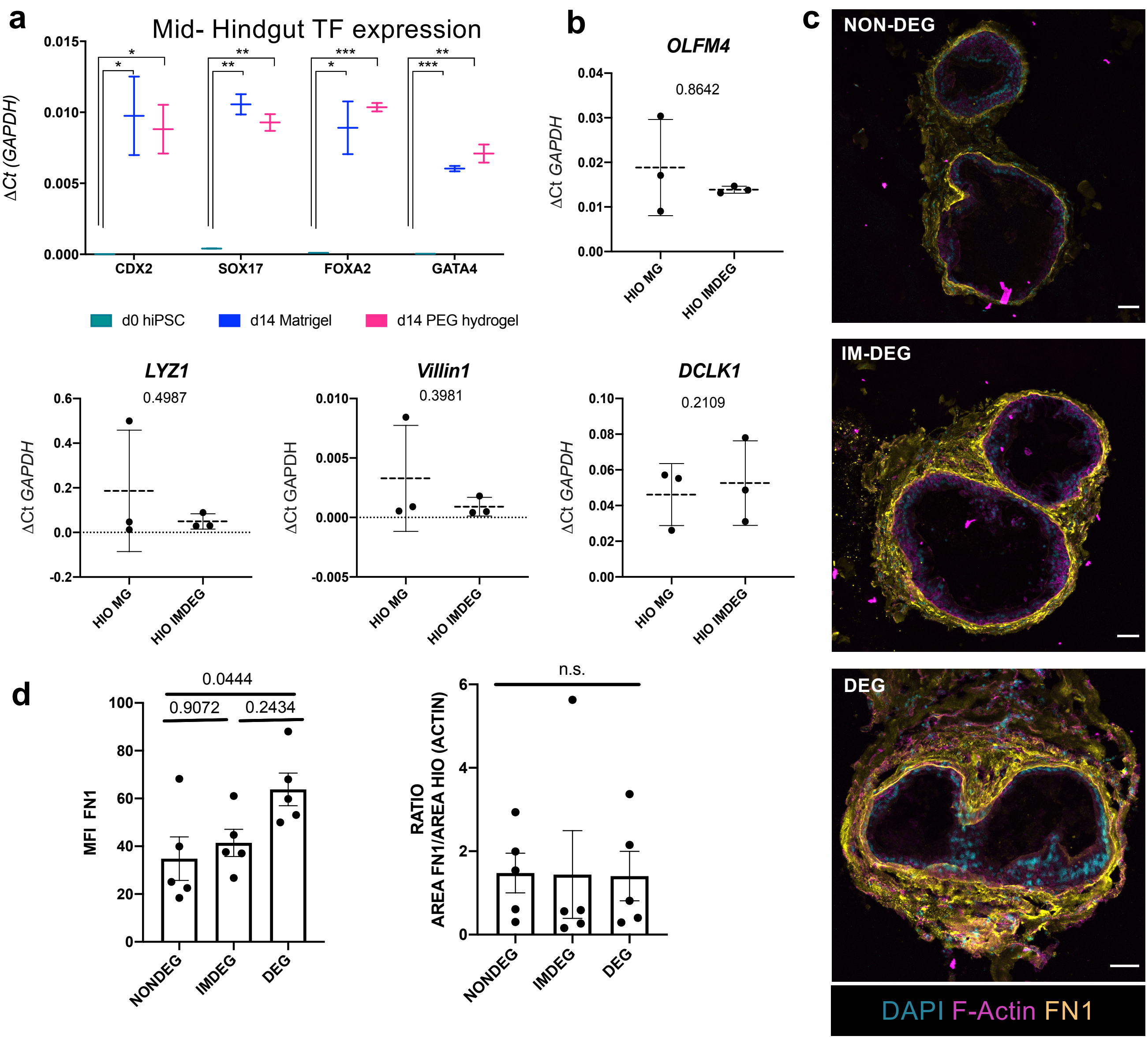


**Supplementary figure 24.** Characterization of human induced pluripotent stem cell (hiPSC)-derived human intestinal organoids (HIOs) in novel synthetic hydrogel system.

a) Expression of endodermal markers (*CDX2*, *SOX17*, *FOXA2* and *GATA4*) is significantly upregulated between day 0 hiPSC and d14 differentiated immature HIO, but not different between HIOs cultured in Matrigel versus IM-DEG PEG hydrogel at day 14, suggesting maturation into hindgut endoderm organoid fate is not adversely affected by PEG hydrogel culture. b) Relative expression of mature gut markers in d75 HIO cultured in Matrigel or post 7-day encapsulation in IM-DEG hydrogel show no significant differences in expression of *OLFM4* (stem cell), *LYZ1 (Paneth Cell), VILLIN1 (Enterocyte), or DCLK1 (Tuft cell)*. c) Representative confocal images of FN1 deposition in DEG, IM-DEG, and NON-DEG hydrogels show that HIO-fibroblasts retain the ability to deposit ECM (Fibronectin1) in the synthetic hydrogel system, making it an appropriate model for further studies. Scale bars 50um, z-projection of 10 stacks. d) FIJI-quantification of confocal images in (c). While more FN1 is deposited in 75% DEG hydrogels than in 0% NON-DEG hydrogels, the area of deposited matrix is not significantly different between conditions (One way ANOVA with Tukey's test.)

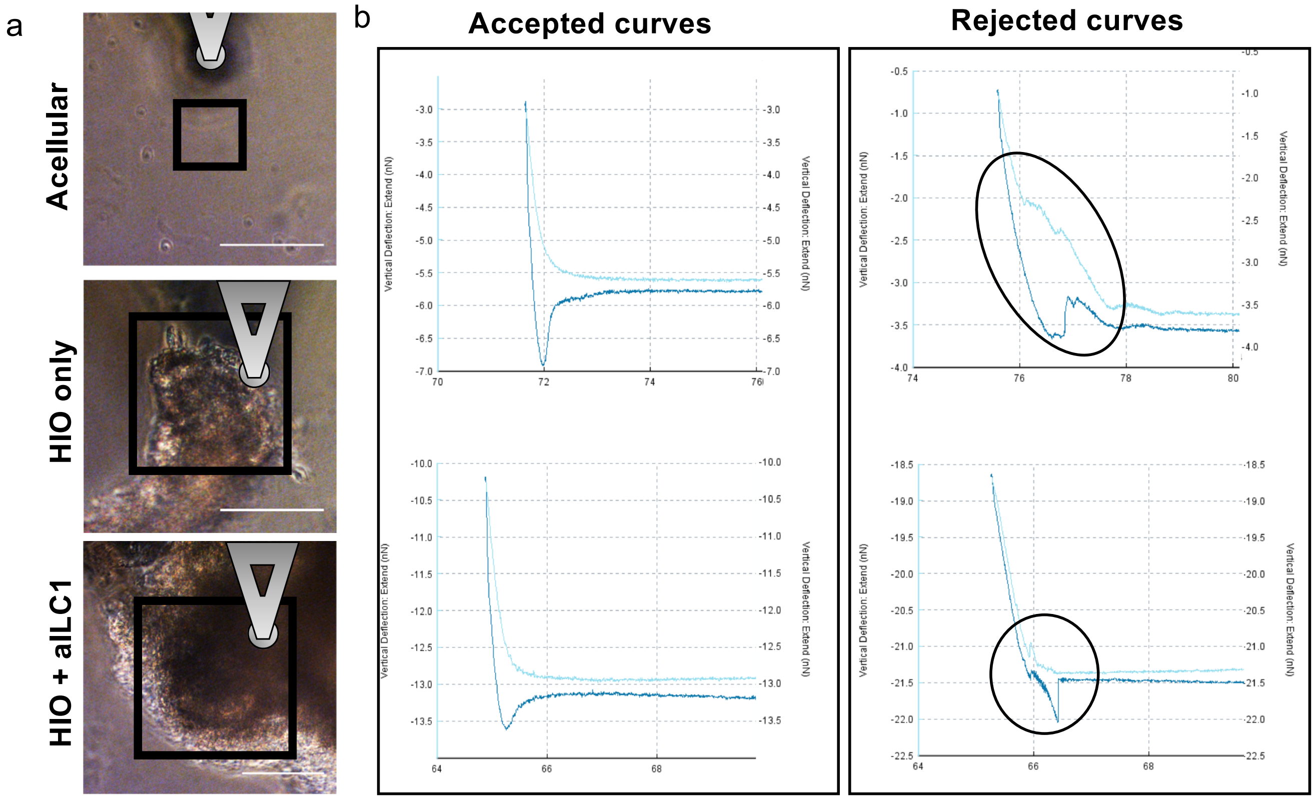


**Supplementary figure 25.** AFM force maps collected on HIO-laden hydrogels

a) Force-distance curves were collected in maps on gels with encapsulated HIO with or without aILC1, with HIO regions determined through morphology in bright field images. Scale bar =100$\mu$m. Boxes highlight areas mapped in examples shown in Fig. 5b (HIO only/HIO+aILC1), and overlaid cartoons show the position of the cantilever. This experimental approach combined with the Hertzian model allows for the measurement of relative changes in $E$ produced by experimental conditions. This is because although the AFM probe (bead d=50$\mu$m) only indents a small distance into the hydrogel surface, underlying soft/stiff hydrogel/secreted matrix will contribute to the material's mechanical response, akin to a "two-spring system", the concept of which is well described in the field of mechanics. In short, when a mechanical load is applied to two springs, with spring constants k1 and k2 in series, the mechanical response of the system will be a combination of both springs' mechanical properties. Here, k1 and k2 can be considered akin to the mechanical properties of the overlying hydrogel and underlying degraded hydrogel/secreted matrix. Such effects have been experimentally verified using AFM-based force spectroscopy measurements on microglial cells cultured on soft polyacrylamide substrates, whereby indentation measurements collected on the overlying cells were impacted by deformation of the underlying soft substrate, which impacted calculations of $E$. (Rheinlaender *et al., bioRxiv* (2019)[1])
b) Example force-distance curves collected during AFM-based stiffness mapping. Force curves were accepted if they had both a smooth clear indentation and a flat baseline (left). Curves for which it was apparent that the indentation had been disrupted (right) were rejected and are presented as crosses (X) on the stiffness maps.

[1]https://doi.org/10.1101/829614

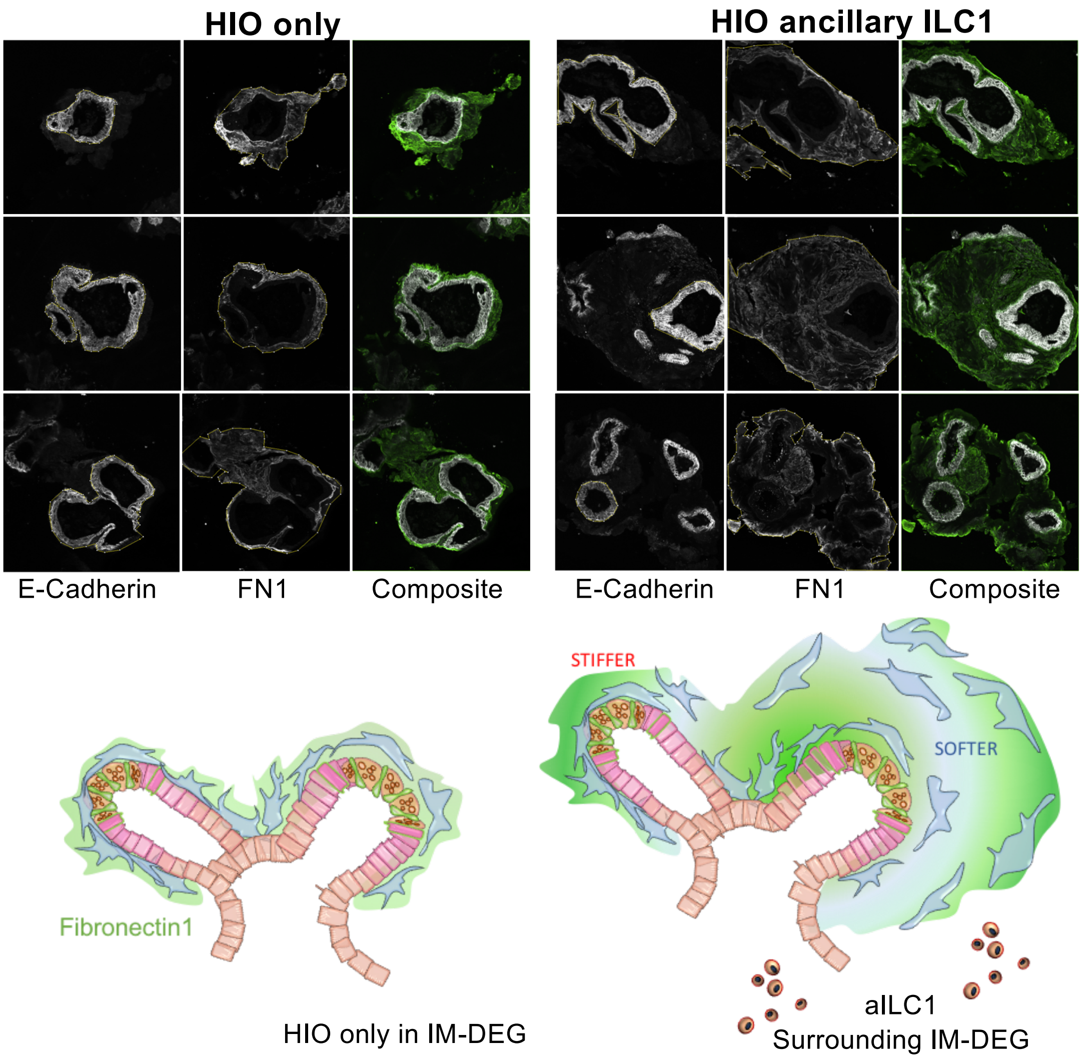


**Supplementary figure 26.** Quantification of fibronectin deposition area

a. Additional representative images of HIO in IM-DEG hydrogels after 4day co-culture either with (right) or without (left) ancillary ILC1. After AFM measurements were taken, HIO gels were PFA fixed and cryosectioned (OCT, 14$\mu$m slices) and co-stained for E-cadherin (white) and Fibronectin1 (FN1, green). The outline of the E-CAD+ epithelial organoid and the edge of the FN1+ remodeled area (green) were detected by binary mapping in FIJI, and the FN1+ area was subtracted from the E-CAD+ area to normalize for the size of the epithelial organoid (N=5 with N=7-10 organoids per condition) to show the relative area of ECM remodelling (relates to Fig. 5g)

Diagrams beneath each condition represents the differential FN1 deposition (green) and matrix softening allowing for fibroblast to spread out (blue, softer, aILC1 condition) but also depositing FN1, making fibroblast-dense regions stiffer.
after 7day co-culture in Matrigel.

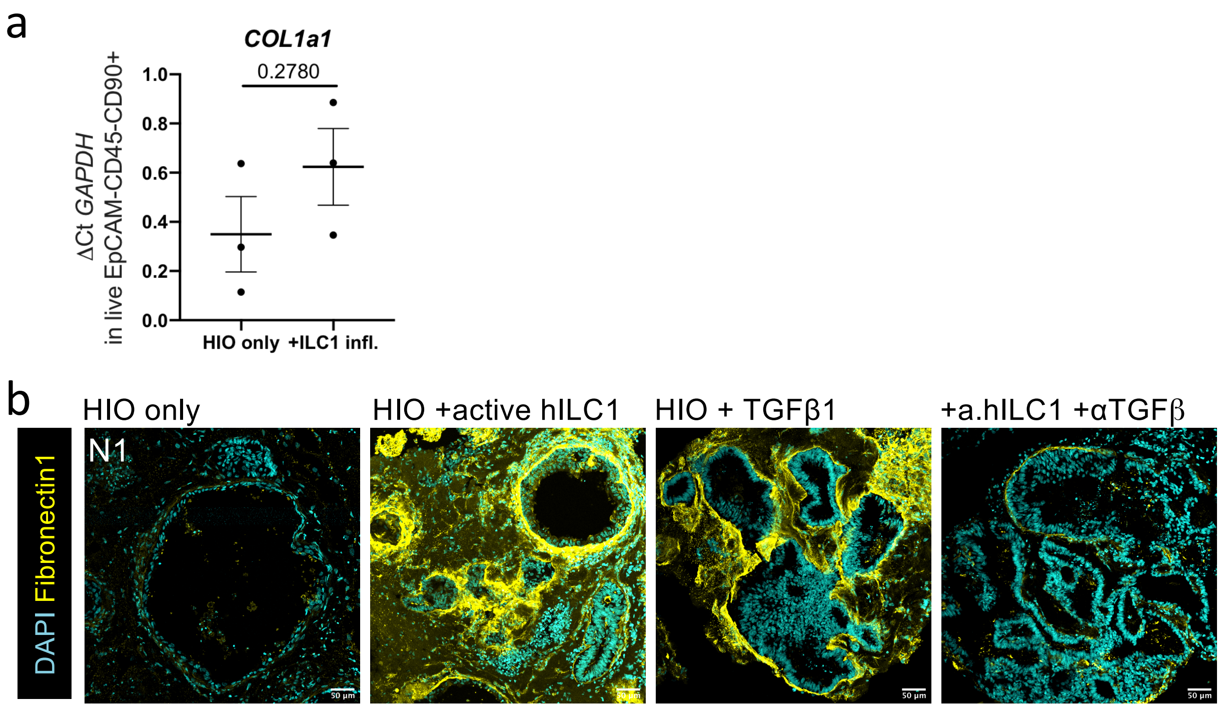


**Supplementary figure 26.** Quantification of fibronectin deposition area

a. RTqPCR of *COL1a1* expression in HIO-derived fibroblasts with or without active hILC1 co-culture. Two-tailed student t-test, error bars represent S.E.M.
b. Representative confocal images of HIO cultured alone, with hILC1, or with addition of recombinant TGFβ1, or with hILC1 and TGFβ1,2,3 neutralizing antibody after 7day co-culture in Matrigel (Rep. of N=2, max projection of 10 stacks in each condition).
